# UNCOVERseq Enables Sensitive and Controlled Gene Editing Off-Target Nomination Across CRISPR-Cas Modalities and Systems

**DOI:** 10.1101/2025.05.09.653165

**Authors:** Kyle J. Kinney, Kun Jia, He Zhang, Ellen Schmaljohn, Thomas Osborne, Bernice Thommandru, Karthik Murugan, Andrea Sánchez-Peña, Sean West, Shengyao Chen, Roshani Codipilly, Morgan Sturgeon, Rolf Turk, Matthew McNeill, Mark Behlke, Ashley Jacobi, M. Kyle Cromer, Garrett Rettig, Gavin Kurgan

## Abstract

The rapid development of CRISPR-Cas gene editing technologies has revolutionized genetic medicine, offering unprecedented precision and potential for treating a wide array of genetic disorders. However, assessing the risks of unintended gene editing effects remains critical, and is complicated by new editing modalities and unclear analytical guidelines. We present UNCOVERseq (Unbiased Nomination of CRISPR Off-target Variants using Enhanced RhPCR), an improved *in cellulo* off-target nomination workflow designed to sensitively nominate off-target sites (<0.01% editing) with defined input requirements and analytical process controls to provide empirical performance evidence across diverse circumstances. Using this workflow, we nominated off-targets across 192 guide RNAs (gRNAs) and demonstrated superior performance compared to existing methodologies. We identified a subset of six gRNAs with a dynamic range of specificity and confirmed the relevance and high true positive rate of our nomination method, providing relative risk assessments for multiple modalities (*S.p.* Cas9 and derived high-fidelity variants / base editors) in a translational system involving hematopoietic stem and progenitor cells (HSPCs). Additionally, we established that double-strand break (DSB) editing retains a strong, positive rank correlation to single-strand break (SSB)-mediated base editing, highlighting the importance of DSB nomination sites as candidate loci for base editing. Overall, UNCOVERseq improves informed risk assessment of gene editing in translational systems by enhancing the quality of off-target nomination.

## Introduction

CRISPR-Cas systems have revolutionized genetic engineering by enabling precise genome modifications. Derived from a bacterial adaptive immune mechanism, CRISPR-Cas9 allows targeted DNA edits in various organisms, enabling vast potential in medicine, agriculture, and biotechnology^1,2^. However, CRISPR-Cas9 faces challenges, notably off-target effects where unintended genomic regions similar to the target sequence can be edited^3,4^. This off-target activity is often due to sequence homology between the protospacer region of the gRNA and non-target regions of the genome. These off-target edits can lead to unwanted genetic changes and disrupt the function of essential genes leading to adverse effects on cellular function, viability, development of secondary diseases, or exacerbate existing conditions that pose risks in therapeutic applications.

Recent advancements in gene editing have driven the parallel development of technologies to nominate off-target editing effects^5–12^. These technologies broadly fall into three categories: cell-based *in cellulo/in situ*, computational *in silico*, and biochemical *in vitro* assays each with distinct advantages and disadvantages. Cell-based assays like GUIDE-seq are known for high precision due to their ability to accurately mimic the cellular environment^7,13,14^. While cell-based methods provide nomination in a cell-type-specific and even donor-specific manner, these methods face challenges in representing diverse genome populations and typically require biologically variable processes such as DNA double-strand breaks (DSBs) and blunt integration of a dsDNA tag through non-homologous end joining (NHEJ) as a mechanism of action. Biochemical *in vitro* assays are highly sensitive and can represent patient genotypes with diverse editing modalities since they are executed on cell-free genomic DNA, but have low precision and suffer difficulties in replicating the cellular environment, which significantly impacts enzymatic activity^6,7,13–15^. Computational *in silico* approaches can rapidly predict all candidate off-target sites with homology to the gRNA in a patient genome / population-specific manner, but result in low precision due to insufficient data to inform predicted enzymatic and biological activity across diverse cell types^11–14,16^. All methods are setback by a lack empirical data and application of standards to characterize sensitivity against known truth, as it is challenging to ascertain a complete list of true positives for each gRNA in an unbiased manner. To better standardize off-target safety assessment, comprehensive information on assessment methods and guidance for their use is essential.

Most methods for off-target nomination currently utilize next-generation sequencing (NGS) to detect off-targets. The efficiency of library preparation, library complexity (i.e., unique genome equivalents sequenced), read depth, and computational analysis methods are all known limitations affecting performance of NGS methods^17^. Some work has endeavored to improve upstream biological and library preparation inefficiencies of *in cellulo* methods^18,19^. Other work has presented evidence of biological optimizations that proxy cell lines promiscuously editing off-targets may represent a superset of off-targets in HSPCs^20^. However, exploration of other NGS variables commonly benchmarked in parallel fields to drive uniformity in experimental procedures and reporting, are still largely lacking for gene editing off-target nomination^17,21^. The FDA has recently worked to address this need through draft guidelines for the cell and gene therapy industry, which currently recommends multiple orthogonal methods for off-target nomination^22^. This guideline for orthogonality ideally aims to mitigate known and unknown weaknesses in nomination, including variability in NGS method execution.

Rapid innovations in gene editing have further complicated the off-target nomination process with the discovery of various editors possessing unique qualities beyond the targeted generation of DSBs with a Type II nuclease. This includes engineered systems generating staggered double-strand breaks (DSBs) in Type V CRISPR-Cas systems or leveraging single-strand DNA breaks (SSBs) coupled to engineered moieties for editing^23–25^. SSB engineered moieties like adenine base editors (ABE)^26^, cytosine base editors (CBE)^27^, and prime editors (PE)^28^ offer benefits such as decreased frequencies of unintended indels and even the ability to make non-indel DNA changes typically reserved for homology-directed repair (HDR). However, they lack consensus guidance regarding appropriate nomination assays for off-target assessment, and previous work has largely used DSB nomination methods as a proxy^29,30^. Due to the nature of these editors to generate both indels and SNPs, an orthogonal approach specializing on each event type may be needed to sensitively nominate off-targets.

In this work, we develop an optimized end-to-end *in cellulo* nomination method and analysis pipeline, UNCOVERseq (Unbiased Nomination of CRISPR Off-target Variants using Enhanced RhPCR), leveraging NHEJ-based dsDNA tag integration for RNase H-dependent PCR (rhPCR)-based library preparation^31^. We demonstrate that using this method with a cell line under promiscuous editing conditions serves as a sensitive proxy for clinically relevant cell types. We establish experimental recommendations for library complexity, coverage depth, and end-to-end process controls that reliably lead to the successful nomination of risk-tiered edited sites with sub-0.01% editing (indel) frequencies observable empirically. By screening 192 gRNAs using UNCOVERseq, we identify an experimental set of gRNAs spanning a broad range of specificities that are broadly representative of use-cases encountered in the field. We find that off-target effects of base editors are directly, rank-order correlated to DSB editing frequencies, demonstrating UNCOVERseq as an effective technology for nominating meaningful off-targets for both current and next-generation editing modalities. We conclude that, regardless of editor modality, the off-target burden of gRNAs quickly decreases to frequencies not measurable – that is, below the limit of detection - with current NGS methods (0.01% for indels and 0.5% for base editing) as gRNA specificity scores rise. However, off-targets may still be observable at higher specificities in an editor/gRNA-dependent manner, emphasizing the need to consistently and empirically nominate and interrogate off-targets across the specificity spectrum.

## Results

### Development and optimization of UNCOVERseq

To create our nomination method, we first started with the original GUIDE-seq protocol and designed a novel orthogonal dsDNA sequence with sufficient length to perform a modified rhPCR to multiplex primers in close proximity within a single reaction while avoiding primer-dimers (Supplementary Table 1). To streamline the process for preparing the nomination gDNA libraries we additionally converted from a mechanical to enzymatic fragmentation (Figure 1A). Upon analyzing data, we observed that freely adaptered dsDNA tag was allocated an average range of 37% to 67% of reads, varying across 4 gRNAs (Figure 1B). This same artifact was also observed with the original GUIDE-seq protocol (Supplementary Figure 1). To improve usable reads resulting from NGS, we introduced a blocking oligo into the PCR1 preparation designed to target the adapter:dsDNA junction (Figure 1A). Introduction of this blocker reduced reads belonging to the adapter:dsDNA artifact to an average range of 0.3% to 0.5%, meaning >99% of reads were now belonging to gDNA:dsDNA junctions (Figure 1B). Nomination frequencies were found to be conserved for all gRNAs (R^2^=0.99) with and without the blocking oligo (Supplementary Figure 2).

**Figure 1.**
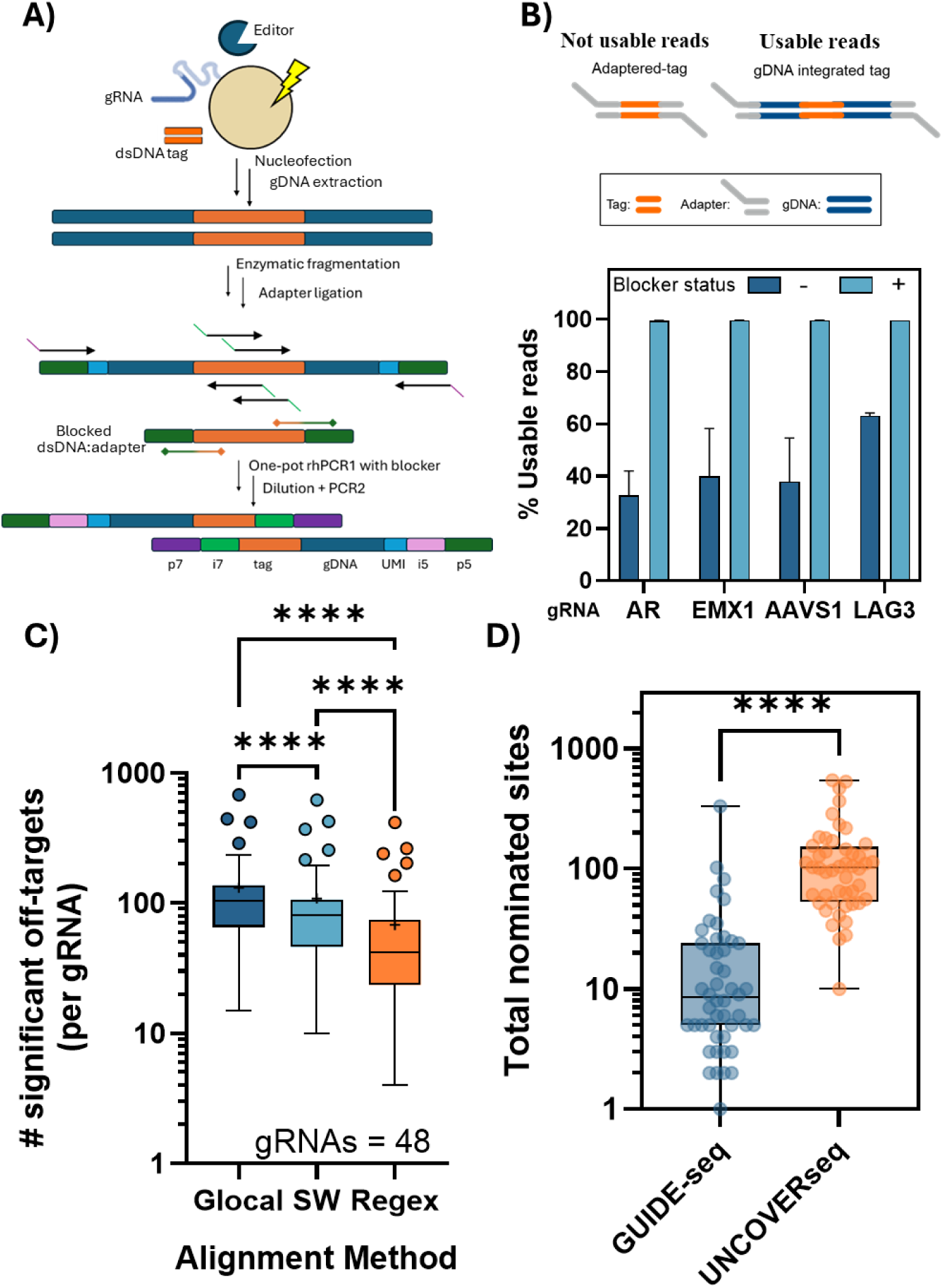
Comparison of UNCOVERseq and the GUIDE-seq off-target nomination workflows. **A)** An overview of the UNCOVERseq workflow demonstrates that cells with a genomically integrated dsDNA tag have gDNA extracted and amplified with rhPCR in a single reaction, with dsDNA tag:adapter byproducts being blocked by a targeted oligo before being sequenced and analyzed using our described workflow. **B)** A depiction is shown regarding what kind of events are targeted by the blocking oligo, with the usable reads (non-dsDNA:adapter reads) measured across nominated off-targets for 4 gRNAs in K562 (n = 3 per gRNA) with (light blue) or without (dark blue) the blocking oligo. **C)** Comparison of different alignment methods used in publicly available nomination packages was performed to determine differences in Levenshtein distance < 7 sites from a glocal Needleman-Wunsch alignment (Glocal; +2/-1/- 10/-1 match/mismatch/gap open/gap extension), Smith-Waterman (SW; original GUIDE-seq pipeline; +2/-1/-100/-100 match/mismatch/gap open/gap extension) or string-match method (Regex; current GUIDE-seq pipeline). **D)** Tukey box plots show the end-to-end differences in total nominated off-target sites for the current GUIDE-seq method (wet lab protocol and Regex alignment) compared with UNCOVERseq. Data is representative of 48 gRNAs (Targets: *PDCD1, LAG3, CTLA4, NRP1, IL2RA*, and *TIGIT*; 8 gRNAs per target). Statistical significance was determined for pairwise comparisons using Wilcoxon matched-pairs signed rank test and for multiple comparisons using a Friedman test with a post-hoc Dunn’s test with Bonferroni correction. ****p < 0.0001.

In parallel to creation of the wet-lab protocol, we created an analysis pipeline with features such as heuristic nomination criteria (Levenshtein distance < 7; read-evidence from both sides of a prospective off-target), statistical comparison of treatment:control samples as nomination criteria (FDR < 0.05) and integrated genomic annotations. We then investigated optimizations in the computational pipeline for nomination of gRNAs using a set of 48 gRNAs spread across the *PDCD1*, *LAG3*, *CTLA4*, *NRP1*, *IL2RA*, and *TIGIT* genes. In off-target nomination, off-target loci are generally determined to be trustworthy based on: 1) frequency, 2) reproducibility, and/or 3) similarity to the intended target sequence (gRNA), with Levenshtein distance > 6 often being used to disqualify an off-target.

To investigate the effect of alignment method used for determination of an off-target list, we tested existing GUIDE-seq pipeline methods (https://github.com/aryeelab/guideseq; commit: 997b892; fuzzy regular expression based; Regex) and historical GUIDE-seq pipeline methods (https://github.com/aryeelab/guideseq; tag:v1.0; Smith-Waterman alignment with -100/-100 gap open/extension penalty) as compared to a glocal implementation of the Needleman-Wunsch algorithm. Investigation of 48 different gRNAs found a significant difference in the number of Levenshtein distance < 7 loci nominated using each approach, and that the glocal Needleman-Wunsch alignment approach yielded a median of 30% and 150% more qualified off-target locations than the current and historical GUIDE-seq analysis approaches (Figure 1C). To do an end-to-end comparison of the current GUIDE-seq method to our method, we did a complete workflow comparison across the same 48 gRNAs and found that our method nominated a range of 60% to 4,150% more off-targets per gRNA, with an average of 132 off-targets as compared to 23 using the GUIDE-seq protocol (Figure 1D). We termed the final instantiation of the end-to-end method as UNCOVERseq (Unbiased Nomination of CRISPR Off-target Variants using Enhanced RhPCR; v1.0).

### Promiscuous cell systems as sensitive UNCOVERseq proxy nomination models

To identify ideal biological operating conditions, we explored biological variables with potential workflow impacts on nomination performance. Promiscuous editing conditions are known to increase editing frequencies at off-targets, which we hypothesize should increase the sensitivity of *in cellulo* methods like UNCOVERseq (Figure 2A). To test this, we selected 4 – 10 gRNAs per cell line and nominated off-targets using UNCOVERseq in K562, iPSCs (wildtype *S.p.* Cas9 or HiFi Cas9), primary T-cells, or a promiscuous HEK293 cell line stably expressing *S.p.* Cas9 (HEK293-Cas9). Investigation of the overlap of off-target frequencies between these cell lines and HEK293-Cas9 found an average of 99.7% to 100% of total UMI-corrected events (corresponding to total frequency) in each cell line could be found in the off-targets of just a single replicate of HEK293-Cas9 (Figure 2B). Comparison of shared nomination frequencies showed a high rank-order correlation for HEK293-Cas9 nominated off-targets between K562 (r = 0.63), iPSC (r = 0.61), and primary T-cells (r = 0.69), demonstrating that the frequency-based importance of different off-targets for prioritization was still largely conserved (Figure 2C-E). We also observed that the overall nominated off-targets number for the same gRNAs could vary significantly in different primary cell lines for the same gRNAs, with low numbers of off-targets nominated in iPSCs, further demonstrating why a promiscuous system can be used to remove this variable (Figure 2F). Overall nomination in a promiscuous cell line like HEK293-Cas9 was capable of generating an average of 196% to 1,560% more candidate targets per gRNA compared to an efficient primary cell type for nomination, like primary T-cells, supporting that this is a more sensitive model for off-target nomination even in translational contexts (Figure 2G).

**Figure 2.**
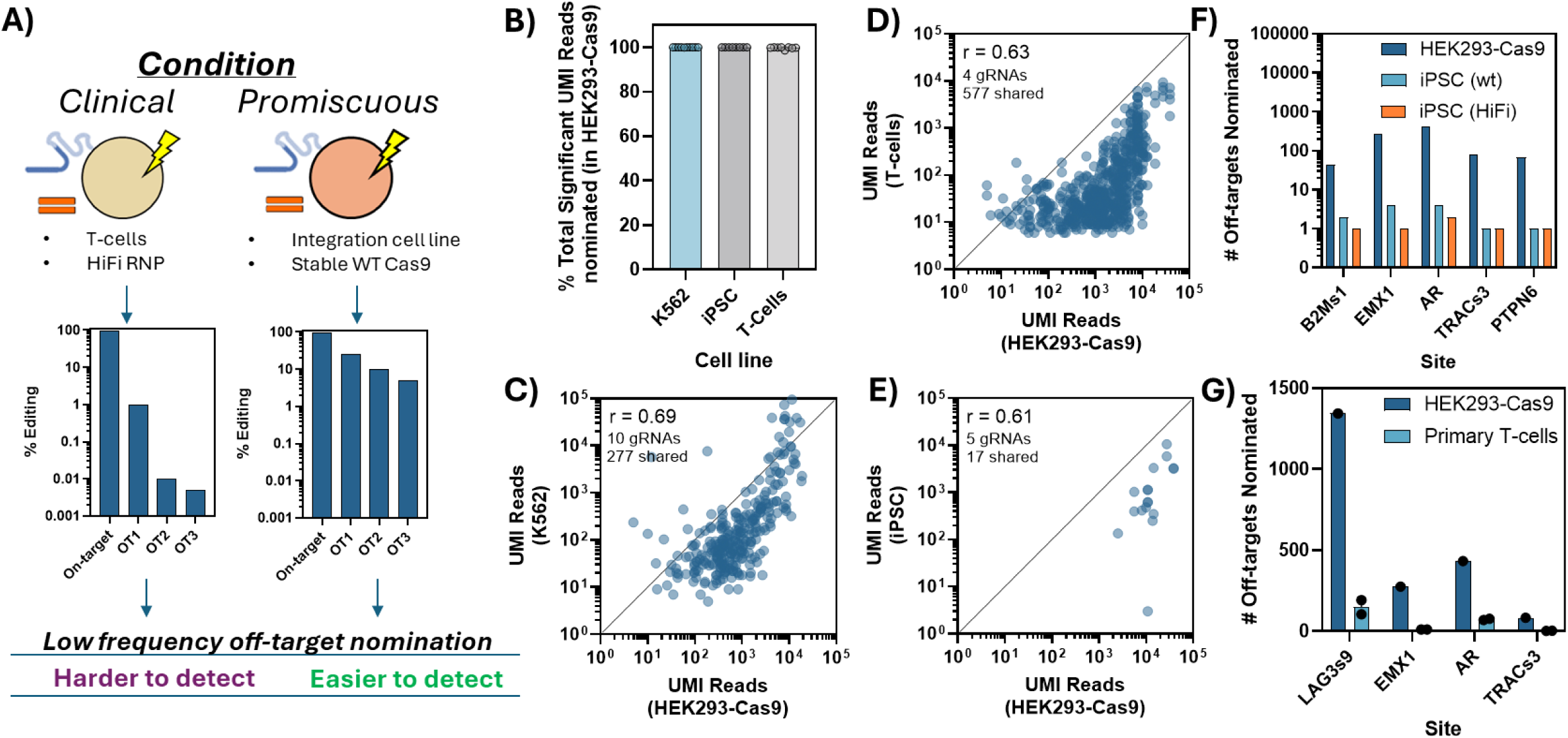
Analysis of performance of a promiscuous cell system and the ability to translate to other cellular systems. **A)** Example diagram demonstrating promiscuous nomination systems can help sensitively represent off-target lists by increasing the frequency that off-targets are detected. **B)** Comparison of total frequencies (represented as the cumulative % significant UMI reads) of nominated sites captured in a single replicate of HEK293-Cas9 vs. wildtype Cas9 RNP transfection using three different cell types: K562 (10 gRNAs; biological duplicates per gRNA), iPSCs (5 gRNAs; one biological replicate per gRNA; wildtype and HiFi Cas9), and primary T-cells (4 gRNAs; 2 biological replicates per gRNA as two different donors). Total UMI-corrected reads, total number of nominated sites, and Spearman correlation are shown for shared nominated sites between HEK293-Cas9 and, **C)** K562, **D)** T-cells, and **E)** iPSCs. Total number of nominated sites between **F)** iPSCs and **G)** primary T-cells are displayed in comparison to HEK293-Cas9.

### Off-target reproducibility using UNCOVERseq

To assess sample-to-sample reproducibility, we compared biological triplicates of UNCOVERseq in HEK293-Cas9 across four gRNAs. An average of 99.2% to 99.7% of instances based on frequency were shared between any two biological replicates, indicating high frequency sites are consistently captured with a single replicate (Figure 3A). Frequency rank order of off-targets was highly conserved (R²=0.997) across replicates as well (Figure 3B). This indicates that both frequencies and sites containing the majority of reads are highly reproducible between UNCOVERseq replicates.

**Figure 3.**
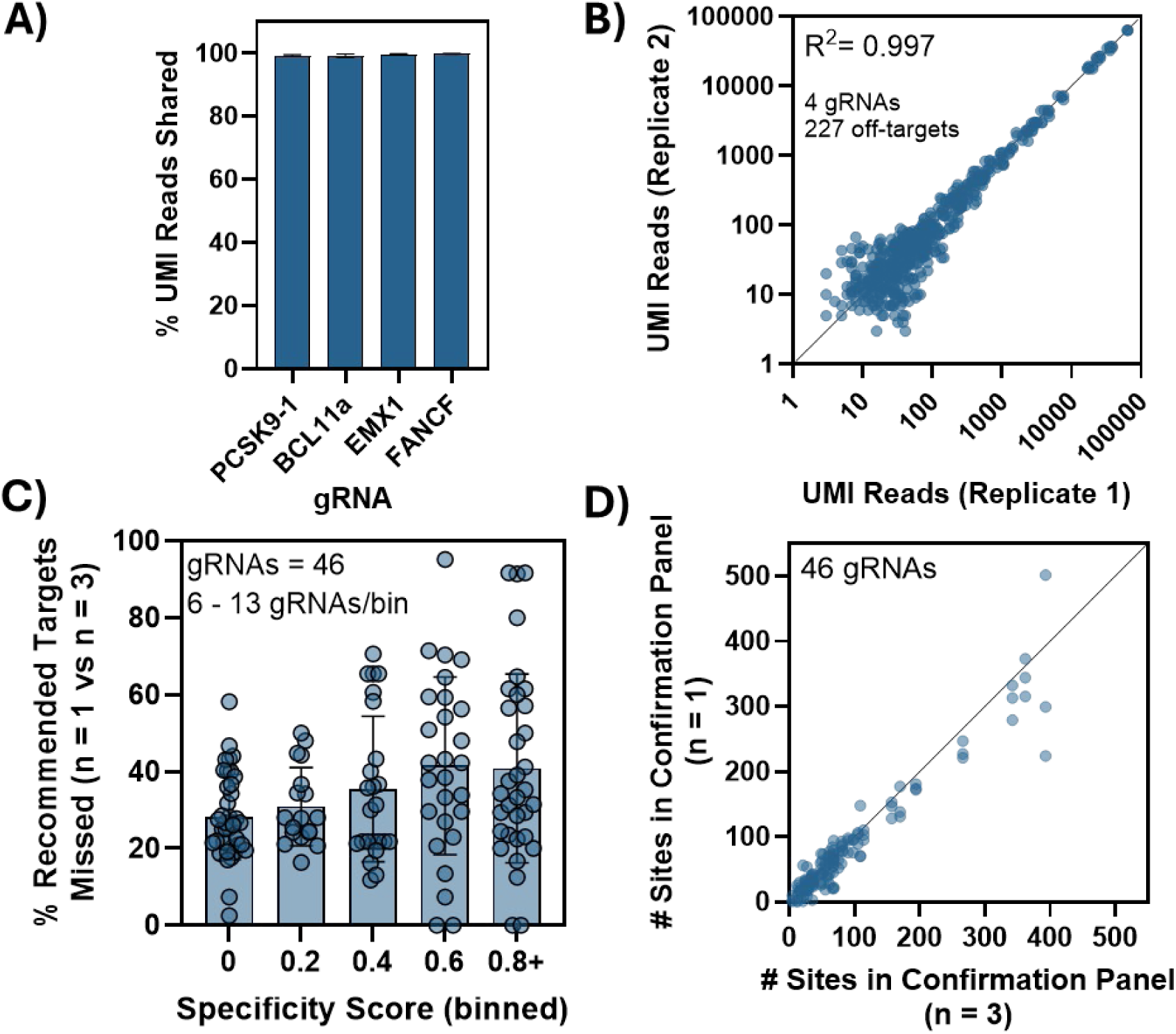
UNCOVERseq nomination reproducibility for high frequency and high priority off-targets. To measure the ability to reproducibly nominate off-targets at consistent frequencies, **A)** a combinatorial comparison of replicates was performed of targets nominated (by frequency, measured as cumulative UMI reads on overlapping targets / UMI reads on all nominated targets) using UNCOVERseq across 4 gRNAs in HEK293-Cas9 (n = 3 biological replicates per gRNA) and **B)** compared the target nomination frequency between replicates. To determine the reproducibility to capture high priority off-targets (defined as Tier 1 to Tier 3 in Supplementary Table 2) 46 gRNAs across a broad specificity score spectrum were nominated in triplicate in HEK293-Cas9 and **C)** the high priority panel content missed and **D)** the total number of off-targets for interrogation are shown for each individual replicate (all nominated sites) compared to the total # of high priority sites (Tier 1 to Tier 3) nominated as a biological triplicate. Specificity score was calculated as the cumulative frequency of UMI-corrected nomination events off-target divided by the number cumulatively between the on-target and off-targets.

To determine ideal experimental conditions for off-target nomination, we characterized factors affecting reproducibility in a functional context. When making decisions about nominated off-targets, ideally off-targets are prioritized based on: a) frequency, b) reproducibility, and c) genomic impact. To this end, we developed a tiering system based on UNCOVERseq data to prioritize off-targets for confirmation (Tier 1 to 3) from less important ones (Supplementary Table 2). To assess reproducibility for prioritizing important off-targets (Tier 1-3), we compared biological triplicates to single replicates of the previous 48 gRNAs in HEK293-Cas9. Without biological triplicates, 30% to 40% of high priority off-targets were not captured or prioritized (Figure 3C). The average frequency of missed high priority targets increased with gRNA specificity, though this is partially because the denominator (total off-targets) is smaller (Figure 2C). While high frequency events were reproducible (Figure 3A), low frequency events lacking full reproducibility or those in important genomic contexts (e.g., exonic) were not appropriately identified without replicates (Figure 3C).

Replication additionally theoretically allows more appropriate tiering of off-targets. To investigate the effect of tiering on the downstream requirements for confirmation, we compared the number of off-targets nominated for appropriately tiered off-targets of biological triplicates (Tier 1 to 3) versus all off-targets of a single replicate. We observe that even though replicates lead to more off-targets nominated, the number of off-targets for interrogation are roughly equivalent when using tiering criteria (Figure 3D). This provides evidence that tiering off-targets in replicate nominations leads to more impactful sites being nominated and recommended for interrogation without significantly increasing the number of off-targets recommended.

### Determination of UNCOVERseq sensitivity and process controls

To characterize the sensitivity of UNCOVERseq and create routine process controls, we extensively characterized the off-targets of a promiscuous gRNA for use as a positive control. Using a promiscuous *LAG3*-targeting gRNA (*LAG3* site 9), we nominated off-targets in 12 biological replicates using HEK293-Cas9. All replicates showed high editing (>90%) with tag integration frequencies between 64% and 81% (Figure 4A). A total of 2269 unique nominated sites were identified, with 723 consistently reproduced across all replicates (Figure 4B). The relative rate at which off-targets could not be reproduced between each subsequent replicate rapidly dropped below <5% after 3 replicates and decreased linearly afterwards (Figure 4B). Although only off-targets reproduced in all replicates were chosen for downstream confirmation, this suggests that after three replicates real off-targets that were harder to reproduce due to low frequencies and random sampling differences may be lost from high replication requirements.

**Figure 4.**
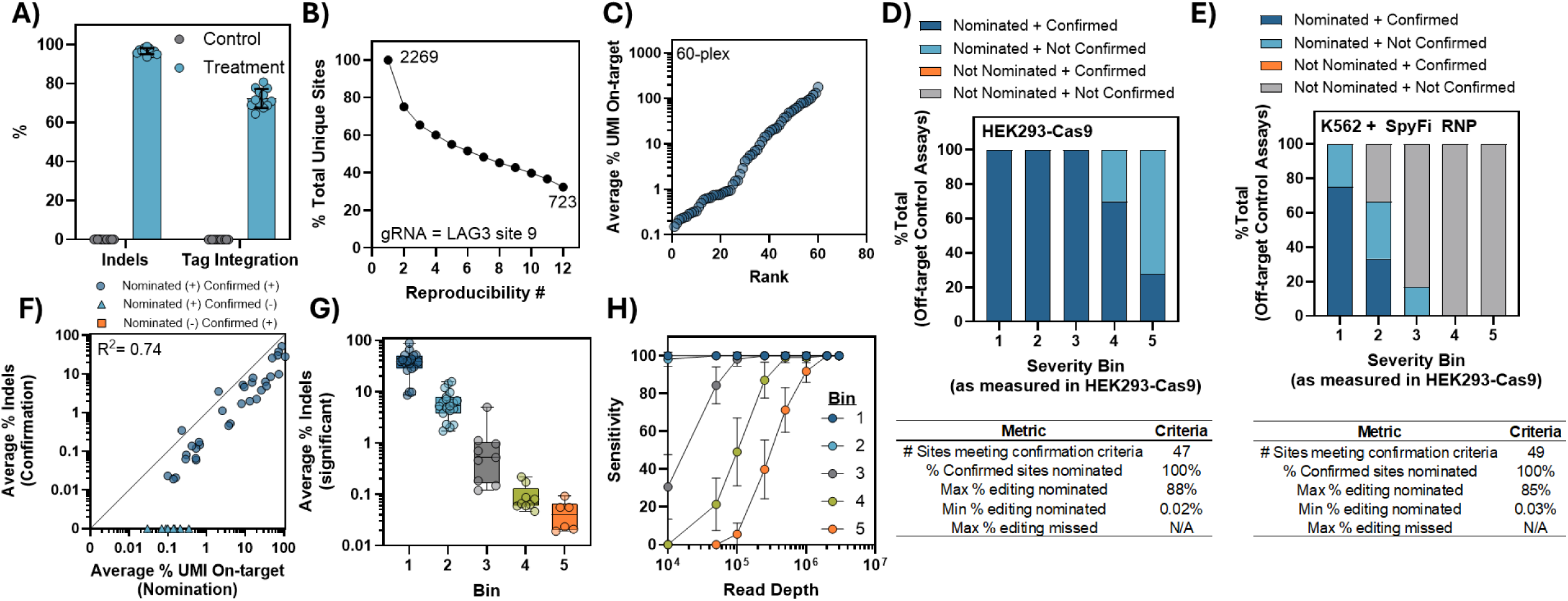
Quality control procedures for confirming assay sensitivity and read requirements. To create a positive control for UNCOVERseq process quality control, we extensively characterized a promiscuous *LAG3* (site 9) gRNA in HEK293-Cas9 using 12 biological replicate transfections with paired controls (no gRNA). Following transfection, **A**) indel frequency and tag integration frequencies were measured at the *LAG3* on-target site via NGS and **B)** the frequency of unique sites relative to the cumulative total reaching each reproducibility frequency is shown. **C)** Twelve sites were selected per frequency bin (Severity Bin 5 = <0.49%; Severity Bin 4 = 0.50% – 0.99%; Severity Bin 3 = 1 - 9.9%; Severity Bin 2 = 10 - 49.9%; Severity Bin 1 = >50% UMI reads relative to the on-target) for routine targeted sequencing with average nomination frequency shown. To test application of this, paired confirmation was performed in **D)** a highly promiscuous condition (HEK293-Cas9; n = 3 biological replicates) and **E)** high**-**specificity condition (K562 nucleofected SpyFi RNP; n = 3 biological replicates) and the status of each of the 60 measured sites meeting coverage criteria (>1,000x) recorded. **F)** Quantification of HEK293-Cas9 confirmation and nomination frequencies is plotted with final status shown as either Nominated and Confirmed (blue circle), Nominated and Not Confirmed (light blue triangle), or Not Nominated and Confirmed (orange square) with **G)** confirmed indel frequencies per bin shown. **H)** Downsampling of the HEK293-Cas9 *LAG3* nomination samples (n = 12 biological replicates) was performed and sensitivity per Severity Bin calculated for recovering all 60 *LAG3* positive control confirmation loci.

To subset off-targets into different groups, we developed five Severity Bins based on the nomination frequency of the off-target relative to the identified on-target event (Supplementary Table 3). These were as follows: Severity Bin 1, >50%; Severity Bin 2, 10 - 50%; Severity Bin 3, 1 - 10%; Severity Bin 4, 0.5 – 1%; Severity Bin 5, <0.5%. To monitor editing frequencies, we selected 12 sites from each frequency bin (Severity Bin), creating a subset of 60 sites for sequencing and confirmation (Figure 4C; Supplementary Figure 3; Supplementary Table 6). Confirmation of editing for these sites in HEK293-Cas9 showed significant indels ranging from 88% to 0.02%, approaching our sensitivity limit (0.01%) (Figure 4D; Figure 4F).

Of the interrogated panel, 100% of sites in Severity Bin 1 to Severity Bin 3 had confirmed editing (Figure 4D). 72.3% of sites in the <0.5% nomination frequency bin (Severity Bin 5) and 30% in the 0.5 to 1% nomination frequency bin (Severity Bin 4) could not be confirmed down to 0.01% indels (Figure 4D). This linear decrease suggests that UNCOVERseq nominates sites with frequencies below 0.01% indels (Figure 4D). This may be due in part to the higher genomic DNA input in UNCOVERseq (500 ng) as compared to the subsequent confirmation library preparation (100 ng). In high specificity conditions (SpyFi Cas9 + ribonucleoprotein nucleofection), fewer off-targets were edited, but a similar dynamic range of confirmed editing was retained (0.03% to 85% indels), with 100% of sites with confirmed editing successfully nominated (Figure 4E). This indicates the approach can be used as a process control to ensure high sensitivity with reportable metrics. UNCOVERseq nomination frequencies were highly correlated (R²=0.74) with confirmation frequencies (Figure 4F). The high true positive rate and consistent decrease in confirmation success in bins approaching our detection limit suggest these sites represent ∼100% true positives.

### Determination of UNCOVERseq input and sequencing requirements

The number of genomes in an amplification reaction and the number of sequencing reads allocated are key limiters for NGS assay performance. To maximize sensitivity, all UNCOVERseq experiments used ∼150,000 genome equivalents. While gDNA input could potentially be increased, we rationalize that this amount of gDNA is attainable by most experimental conditions and represents the ability to potentially detect down to 0.001%, which is below the limit of detection for any currently published confirmation techniques for CRISPR gene editing.

To characterize read depth requirements for reproducible off-target nomination, total reads were downsampled from the previous *LAG3* site 9 UNCOVERseq dataset (n = 12) to frequencies ranging from 3 million to 10,000 reads per sample. Prior to downsampling, editing across confirmable sites showed interquartile range (IQR) frequencies for each Severity Bin as follows: Severity Bin 1, 26%-48%; Severity Bin 2, 2.6%-7.6%; Severity Bin 3, 0.13%-0.82%; Severity Bin 4, 0.06%-0.13%; and Severity Bin 5, 0.01%-0.02% (Figure 4G). Thus, this dataset represents the full dynamic range of what can currently be detected by confirmation (>0.01% indels). High frequency sites in Severity Bin 1 and Severity Bin 2 were nominated with 100% sensitivity using 50,000 reads per sample (Figure 4H). To nominate down to 0.01%-0.02% indel frequencies with 100% sensitivity, at least 2 million reads per sample were required (Figure 4H). Given the tendency of HEK293-Cas9 cells to over-nominate off-targets, we recommend performing UNCOVERseq with >500,000 reads per sample to aim for >50% analytical sensitivity at Severity Bin 5, and >2 million reads per sample for maximum sensitivity in assessing candidate off-target sites (Figure 4H). It is possible that read depth requirements may vary with off-target number. However, by using a promiscuous gRNA (specificity score = 0.013) to determine this value, we propose that this represents the number of reads to successfully nominate sites even with gRNAs with very poor specificity.

### Comparative analysis of UNCOVERseq to other nomination methods

A comparative analysis of UNCOVERseq to peer-reviewed accounts of other nominations methods was performed to better understand how the sensitivity and nomination frequencies of diverse methods compare. Due to variable operational conditions, precision, and total nomination list sizes reported of different methods, we postulate that sensitivity is most appropriately measured using either off-targets with confirmed editing or off-targets from methods with high precision. Interrogation of the 60 *LAG3* site 9 gRNA off-targets compared to CHANGE-seq^7^ and GUIDE-seq^7^ nominations demonstrated that both methods could nominate the most frequent group of confirmed off-targets (Severity Bin 1) with 91% - 100% sensitivity, but sensitivity rapidly decreased in the lower frequency off-target bins. CHANGE-seq was demonstrated to have a sensitivity between 66% to 75% for recovering Severity Bin 3 to 5 off-targets, while GUIDE-seq had a linear decrease from 16% to 0% for these same Severity Bins of confirmable off-targets (Supplementary Figure 4).

Random sampling of the *LAG3* site 9 dataset with 100% reproducibility showed UNCOVERseq-nominated sites had ∼100% precision, with confirmation frequencies correlating to average nomination frequencies (Figure 4). Using this logic, we postulate the full 723 sites in this fully reproducible set are also likely to represent true positives. Investigation of sensitivity and frequencies of previous accounts of CHANGE-seq and GUIDE-seq for this gRNA yielded similar trends in sensitivity per Severity Bin, further supporting this (Figure 5A). This provides additional evidence that reproducible UNCOVERseq sites have high likelihood of being true positives and may serve as an appropriate proxy for measuring sensitivity of different methods.

**Figure 5.**
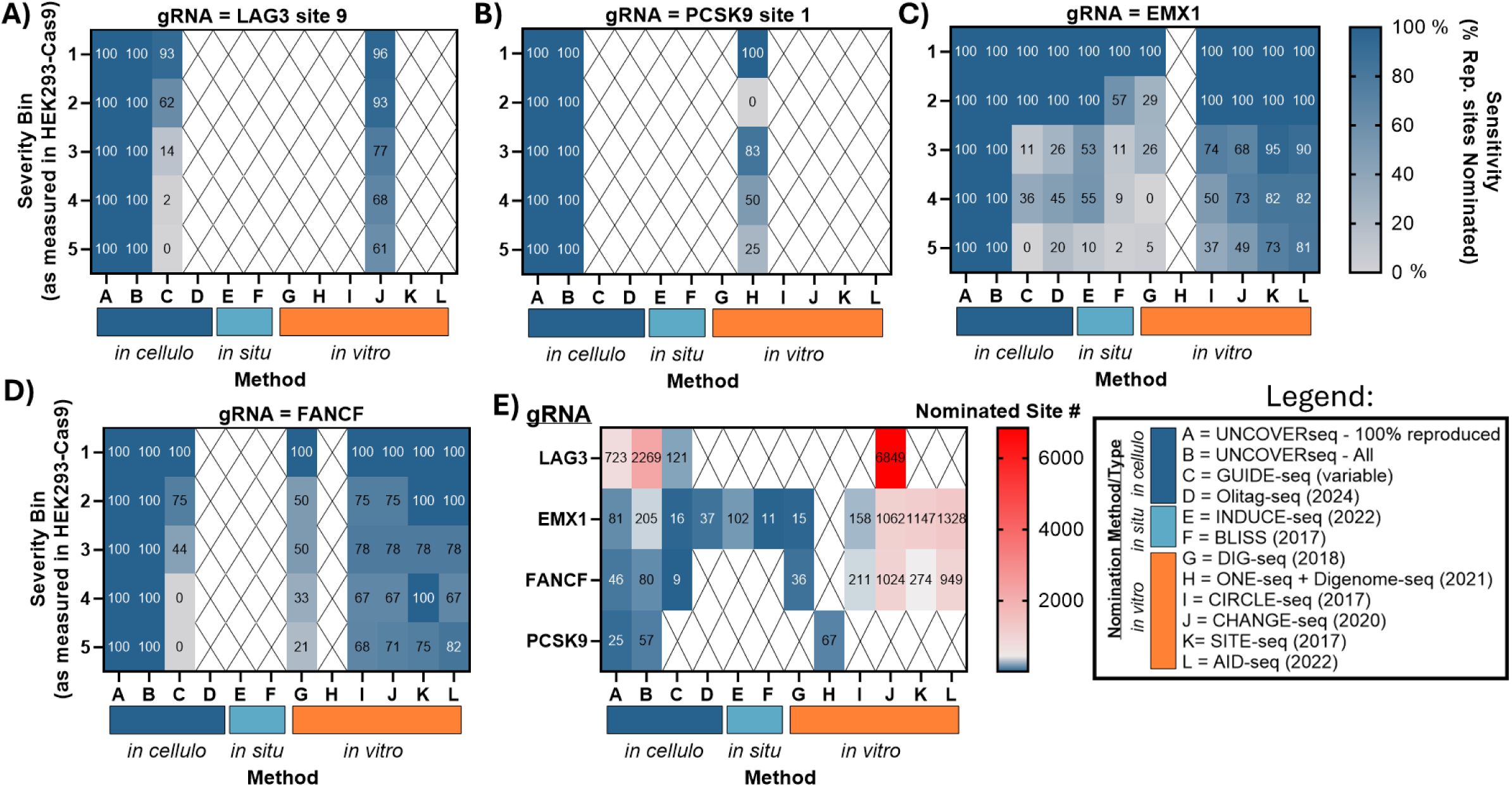
Comparative analysis of UNCOVERseq to other nomination technologies. Comparison of the sensitivity of other published accounts of nomination technologies to nominate 100% reproducible UNCOVERseq off-target sites using **A)** LAG3 site 9 (n = 12 nomination replicates) **B)** PCSK9 site 1 (n = 6 nomination replicates) **C)** EMX1 (n = 6 nomination replicates) and **D)** FANCF (n = 6 nomination replicates). **E)** Total number of off-targets nominated per nomination method per gRNA with color gradient shifts at the number of off-targets easily interrogated using amplicon sequencing (<250 targets; blue/grey) vs those more challenging to interrogate using amplicon sequencing (>250; grey/red). Legend displays the corresponding method to each column in the heatmaps along with the corresponding method type (*in cellulo, in situ, in vitro*). Rep. sites; Reproducible UNCOVERseq sites between all replicates. UNCOVERseq – All represents all Tier 1 to Tier 3 sites from the experiments.

Using our previous finding that off-targets with >3 replicates reproducing a site is likely indicative of true positives (Figure 4B), we compared fully reproduced off-targets from 6 biological replicate UNCOVERseq experiments for three gRNAs (*EMX1*, *FANCF*, and *PCSK9*) to previous accounts of off-targets from GUIDE-seq^5^, INDUCE-seq^10^, Olitag-seq^18^, CHANGE-seq^7^, CIRCLE-seq^6^, AID-seq^32^, BLISS^33^, ONE-seq^34^, DIG-seq^35^, Digenome-seq^34^, and SITE-seq^8^ nomination methods (Supplementary Table 5). For PCSK9, only a combined set of ONE-seq and Digenome-seq sites was available, and nomination sensitivity was found to fluctuate from 0% to 83% through Severity Bins 2 to 5 (Figure 5B). For *EMX1*, DIG-seq and BLISS sensitivity decreased to 29% and 57% after Severity Bin 1, respectively, and sharply decreased at lower frequency bins. Cell-based methods GUIDE-seq, Olitag-seq and INDUCE-seq sensitivity dropped after Bin 2 (11% to 53% sensitivity), ranging from 55% to 0% for off-target nomination frequencies observed after Severity Bin 2 (Figure 5C). However, nomination frequencies for GUIDE-seq, Olitag-seq and INDUCE-seq correlated well with expected frequencies (R^2^ = 0.52 to 0.70), if identified (Supplementary Figure 5). Remaining biochemical *in vitro* nomination methods had the highest sensitivity compared to UNCOVERseq, with approximate rank-order as follows: AID-seq/SITE-seq > CHANGE-seq > CIRCLE-seq (Figure 5C). Sensitivity dropped for all these methods at Severity Bin 3, and ranged from 37% to 95% for Severity Bin 3 to 5 (Figure 5C). For *FANCF*, GUIDE-seq and DIG-seq sensitivity quickly decreased below 50% after Severity Bin 2 to a range from 0% to 21% below Severity Bin 3 (Figure 5D). AID-seq/SITE-seq achieved higher sensitivity compared to CIRCLE-seq/CHANGE-seq, similar to previous observations with EMX1 (Figure 5C-D). CHANGE-seq and CIRCLE-seq sensitivity dropped to a range of 67% to 78% after Severity Bin 1 (Figure 5D). SITE-seq and AID-seq sensitivity dropped after Severity Bin 2 and ranged from 75% to 100% (Figure 5D).

In this calculation of sensitivity per method, each method nominated a widely variable number of sites per gRNA, which affects this calculation. The number of nominated sites ranged from 9 for *FANCF* using GUIDE-seq to 6,849 off-targets using *LAG3* site 9 with CHANGE-seq (Figure 5E). While biochemical *in vitro* methods like CHANGE-seq, AID-seq, and SITE-seq showed the greatest comparative sensitivity, they also had the largest nominated site list, ranging from 291 to 1,328 sites for the *FANCF* and *EMX1* gRNAs shared between most methods (Figure 5E). Investigation of method normalized nomination frequencies demonstrate that *in vitro* methods have the poorest correlation to UNCOVERseq derived frqeuencies (R^2^ = 0.01 to 0.26), with AID-seq and SITE-seq having the lowest correlations (Supplementary Figure 5). Nomination frequencies derived from cell-based *in cellulo* and *in situ* methods (GUIDE-seq, OliTag-seq, INDUCE-seq, BLISS) better correlate to UNCOVERseq nomination frequencies (R^2^ = 0.31 to 0.70), which previously were shown to correlate to observed confirmation frequencies (Supplementary Figure 5; Figure 4F; Figure 5). These findings demonstrate that UNCOVERseq improves upon the sensitivity of existing *in cellulo* methods such as GUIDE-seq, in addition to subsequent improved methodologies such as OliTag-seq. Our findings also demonstrate that *in vitro* methods are not inherently more sensitive than *in cellulo* methods for discovering true off-targets, and UNCOVERseq nominates confirmable off-targets not detected in other methods.

### Screening gRNAs of variable specificity

To identify the specificity of a broad set of gRNAs for future experimental design, 192 gRNAs were selected and UNCOVERseq performed in HEK293-Cas9 cells. Samples were sequenced to a median of 1.8 million reads, in line with our previous recommendations for maximizing sensitivity (Supplementary Figure 6A). Following this, we calculated a specificity score for each gRNA as previously defined^7^. We rationalized that only a single replicate per gRNA was needed for this experimentation since we were most interested in higher frequency off-targets that would be recovered from a single replicate in the promiscuous system (Figure 3A). A range of specificity scores per gRNA were recovered ranging from 0.00063 to 1.0 (Supplementary Figure 6B). Comparison to previously published CHANGE-seq specificity scores of overlapping gRNAs demonstrated a high rank-order correlation (r = 0.63) between specificity scores derived from both methods, though sometimes large disagreements in specificity scores for higher specificity gRNAs occurred (Supplementary Figure 6C). We then subset gRNAs by specificity scores (binned in increments of 0.2) and obtained a relatively uniform distribution with each bin containing a range between 22 to 47 gRNAs each (Supplementary Figure 6D-E). We further subset these gRNAs by those that were ABE- and CBE-compatible as defined as having either an “A” in the +4 to +7 positions (5’ to 3’) or a “C” in the +4 to +8 positions (5’ to 3’). This resulted in a less uniform distribution of gRNAs across the specificity spectrum with a range of 8 to 21 gRNAs per specificity score bin (Supplementary Figure 3D-E). From this we selected six gRNAs for further experimentation as representing a continuous range from 0 to 1 supporting all editor modalities: *PDCD1* site 8, *CYP2C18*, *RNF2*, *TRAC* site 7, *B2M* site 1 and *TIGIT* site 7 (Figure 6A).

**Figure 6.**
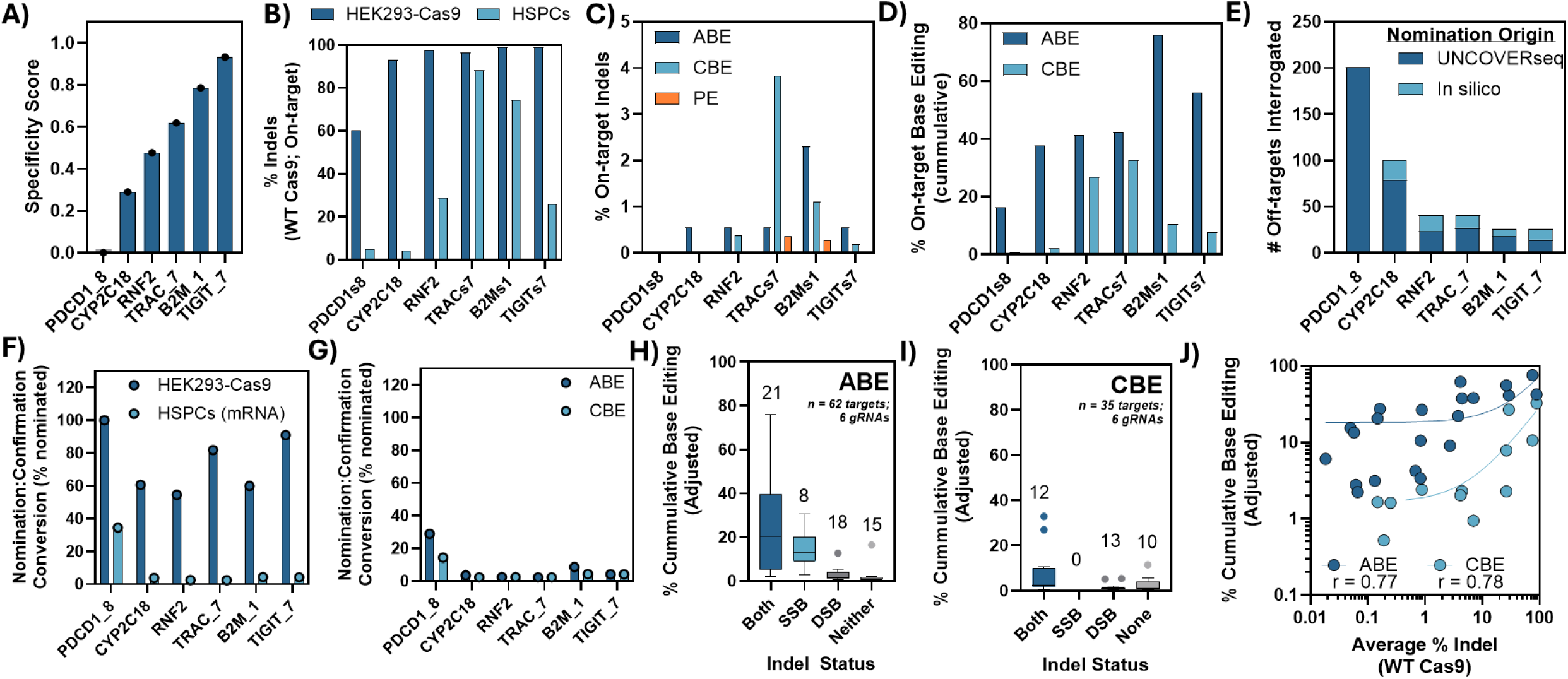
On-target and off-target editing in HEK293-Cas9 and HSPCs. **A)** Six gRNAs with a single targeted ABE or CBE base were selected with specificity scores shown. HEK293-Cas9 (n = 1) and HSPCs (n = 3 donors), with simultaneous delivery of the corresponding HiFi Cas9 mRNA, were delivered each gRNA and had **B)** on-target indel editing quantified by NGS. HSPCs were also delivered mRNA with either the *S.p.* Cas9-ABE8 or *S.p.* Cas9-CBE fusion for each gRNA and had **C)** indel editing and **D)** base editing quantified using NGS. **E)** Multiplexed amplicon sequencing (rhAmpSeq) panels were created for each gRNA for off-target quantification based on origin from UNCOVERseq or *in silico* nomination (*in silico)* with the number of targets interrogated per gRNA shown. After sequencing/confirmation of off-targets in all conditions, the frequency that UNCOVERseq nominated sites converted to true positives was measured for **F)** *S. p.* Cas9 or HiFi Cas9 indel editing and **G)** *S. p.* Cas9 base editing conditions. Confirmed base editing sites were categorized by their respective indel confirmation status (DSB = wildtype *S.p. Cas9;* SSB = ABE/CBE nickase) with cumulative base editing plotted for **H)** ABE and **I)** CBE editing conditions. **J)** Frequencies were plotted for all confirmed ABE/CBE sites with both SSB and DSB confirmation with the respective DSB indel frequency and Spearman r calculated. Line indicates linear regression for ABE (dark blue) and CBE (light blue).

### Comparative analysis of editors in HEK293-Cas9 and HSPCs (on-target)

Next, we sought to determine the translation of UNCOVERseq off-targets across a broad range of specificities in a translational ex-vivo system (HSPCs with mRNA editor nucleofection) across different editing modalities (DSB Cas9, SSB Base Editors, SSB Prime Editors). To do this, we edited HSPCs with one of six gRNAs along with mRNA of either a) a high fidelity *S.p.* Cas9,variant (HiFi Cas9) b) *S.p.* Cas9 (D10A) fused to a Cytosine Base Editor (AncBE4max^36^), c) *S.p.* Cas9 (D10A) fused to Adenine Base Editor (ABE8e^37^) and d), *S.p.* Cas9 (H840A) fused to the PE2 system^28^ with a pegRNA intended to introduce a single SNP. HEK293-Cas9 was also edited in parallel with the wildtype *S.p.* Cas9 nuclease. Evaluation of on-target *S.p.* Cas9 editing found that editing in HEK293-Cas9 was highly efficient at all sites, ranging from 60.4% to 99.4% indel editing, but with a trend of decreased frequencies at lower specificity gRNAs (Figure 6B). Evaluation of on-target indel editing in HSPCs had a range of 4.4% to 88.4% indel editing for the DSB-based HiFi Cas9 editor, 0.0% to 2.3% indel editing for ABE editors, and 0.0% to 3.8% indel editing for CBE editors, and 0.0% to 0.36% indel editing for Prime Editors (Figure 6C). Intended on-target cumulative base editing ranged from 16.5% to 75.9% for ABEs, and from 0.78% to 32.9% for CBEs (Figure 6D). No significant base editing was observed for either the DSB-based *S.p.* Cas9 editors or SSB prime editor, as would be expected (Data not shown). No significant frequencies were observed for the intended mutation to be introduced via prime editing at any sites (Data not shown). For this reason, we excluded further evaluation of the prime editor. Since we show evidence of on-target indel editing with the PE construct, we hypothesize the lack of intended activity is due to the need for substantial pegRNA optimization to achieve successful prime editing, as has been previously reported^38^. For all editors in HSPCs, similar trends were observed to HEK293-Cas9 that the lower specificity gRNAs *PDCD1* site 8 and *CYP2C18* had trends with decreased editing, suggesting that these sites may be overall less effective at the on-target, potentially due to competing off-targets (Figure 6). We conclude from this that on-target editing was successful in all conditions except prime editing, with highly variable frequencies as may be expected without substantial optimization.

### Comparative analysis of DSB editors in HEK293-Cas9 and HSPCs (off-target)

A range of nominated off-targets for the six gRNAs were selected from two orthogonal methods for downstream confirmation: UNCOVERseq nominations and *in silico* nominations. A range of 26 to 201 putative editing sites were interrogated per multiplexed amplicon rhAmpSeq panel with an UNCOVERseq:*in silico* nomination origin split ranging from 53.8% to 100% for the interrogated target lists (Figure 6E). Panels were sequenced with a goal of at least reaching 1,000x coverage per off-target to ensure adequate sensitivity for calling significantly edited off-targets. Sequencing the six panels demonstrated we were able to achieve a median read coverage ranging between 34,000x to 76,000x, with consistency in coverage between edited and corresponding control samples (Supplementary Figure 7A). The number of off-targets reaching sufficient coverage (>1,000x) ranged from 92% to 100% per panel, with a range of 0 to 15 targets not being at sufficient coverage per panel (Supplementary Figure 7B).

To determine the frequency of UNCOVERseq HEK293-Cas9 nominations that convert to empirically edited sites in variable DSB editing contexts, we compared this frequency for *S.p.* Cas9 both in HEK293-Cas9 and HSPCs (HiFi Cas9). For HEK293-Cas9, a range of 54.5% of nominated off-targets all the way to 100% of off-targets had confirmed editing ranging from to 0.02% to 95% indel editing, demonstrating the true positive rate for UNCOVERseq-nominated sites remains high even with only a single replicate in the appropriately paired confirmation context (Figure 6F; Supplementary Figure 8A-F). For HSPCs, a range of 2.4% to 34.5% of nominated targets were successfully confirmed per gRNA, with confirmed indel editing ranging from 0.06% to 88% (Figure 6F; Supplementary Figure 9A-F). Nomination:confirmation frequencies trended to increase as gRNA specificity decreased, suggesting that the method is still successfully nominating relevant off-targets, but that these sites likely no longer exceed detectable frequencies or are no longer edited in the higher genome editing specificity context of HSPCs delivered a HiFi Cas9 mRNA editor (Figure 6F). Furthermore, at higher gRNA specificities the only nominated target being confirmed is the on-target site for HiFi Cas9 in HSPCs (Supplementary Figure 9A-F).

Off-targets that were confirmed were compared to the list of those that would have been dropped given a different previously evaluated alignment method (Regex method; Figure 1). A range of 3 to 23 bona fide off-targets per gRNA in HEK293-Cas9 were successfully nominated using our alignment method that were missed using GUIDE-seq analysis Regex method, with a range of observed indel editing from 0.02% to 68% (Supplementary Figure 10A). In HSPCs, 1 bona fide HiFi Cas9 off-target was identified with a frequency of 0.01% with our alignment method that was missed using the Regex alignment method (Supplementary Figure 10B). This demonstrates that DSB off-target sites that were nominated due to differences in alignment criteria can result in bona fide off-target indel editing in both HEK293-Cas9 and HSPCs.

### Comparative analysis of non-DSB editors in HSPCs (off-target)

Off-target were simultaneously confirmed for both indel and base editing in the non-DSB treatments for HSPCs (ABE and CBE). Similarly, we interrogated the frequency that UNCOVERseq HEK293-Cas9 nominations convert to empirically edited sites in HSPCs being delivered a base editor. For ABE treatments, a range of 2.4% to 29% of nominated targets had confirmed editing ranging from 0.53% to 75.9% cumulative ABE editing (Figure 6G-H; Supplementary Figure 11A-F). For CBE treatments, a range of 2.4% of nominated targets to 14.5% of targets had confirmed editing ranging from 0.51% to 32.9% CBE editing (Figure 6G; Figure 6I; Supplementary Figure 12A-F). Significant indel editing was observed for all ABE and CBE editing treatments, with largely only the on-target gRNA containing indels at higher specificity gRNAs (Figure 6C; Supplementary Figure 13A-F; Supplementary Figure 14A-F). Off-target indel frequencies for ABE treatments ranged from 0.02% to 0.88% indels across different gRNAs (Supplementary Figure 13A-F). Interestingly, three off-target sites were found to generate indel events at the higher specificity *TRAC7* gRNA under ABE treatment conditions, which lacked any significant off-targets in paired DSB Cas9 treatment (Supplementary Figure 13D). Off-target indel frequencies for CBE treatments ranged from 0.08% to 0.66% indels across different gRNAs (Supplementary Figure 14A-F). Generally, it was observed that indel and base editing frequencies were lower in CBE-treated samples in comparison to ABE-treated samples (Figure 6I-J; Supplementary Figure 11A-F; Supplementary Figure 12A-F), although this could be a result of lower overall activity of the base editor instead of off-target propensity.

When comparing the list of confirmed ABE/CBE off-targets to those that would have been excluded given a different alignment method during nomination we found 1 bona fide off-target of the *PDCD1* gRNA that was identified for both ABE and CBE treatments with a frequency range of 0.5% to 3.1% base editing that was missed using the Regex alignment method (Supplementary Figure 10B). This demonstrates that off-target sites that were nominated due to differences in alignment criteria can also result in bona fide off-target base editing activity for both ABE and CBE editors in HSPCs.

To investigate relationships between DSB indels, SSB indels, and base editing, we binned confirmed base editing off-targets based on their presence of indels in either DSB or SSB systems. Base editing with the highest frequencies (median 20.6% and 2.3% for ABE and CBE, respectively), were found to coincide with indel editing for both DSB and SSB systems (Figure 6H). Interestingly, only ABE treatments were found to have an increased frequency of base editing at SSB only sites, with eight detected SSB-only off-targets with a median 13.1% cumulative base editing compared to zero sites for CBE (Figure 6). This may coincide to activity differences, as both ABE editing/indel activity was generally higher than CBE editing/indel activity across the different sites (Supplementary Figure 11-14). DSB-only and sites with no evidence of significant indel editing were present in confirmed sites for both ABE and CBE treatments, albeit with lower median cumulative base editing frequencies (Figure 6). The on-target indel and base editing activity of the different gRNAs were rank-order correlated (r=0.66 - 0.89), suggesting that indel editing frequencies may be predictive of base editing frequencies (Supplementary Figure 15). Similarly, off-target DSB indel editing frequencies from HiFi Cas9 demonstrated rank-order correlation with off-target base editing frequencies (r = 0.77 – 0.78) at sites that had significant DSB indel editing frequencies in HSPCs (Figure 6J). This provides evidence that DSB editing may be indicative of base editing activity, meaning that DSB-nominated sites are meaningful for interrogation in the context of both indel and base editing off-target assessment for both ABE and CBE modalities.

### Comparative translocation analysis and overall editing burden across editing modalities

To investigate differential frequencies of editor modalities to generate large structural variants (>0.1% frequencies) in HSPCs, we investigated the previously described six sites for on-target:off-target and off-target:off-target translocations using amplicon sequencing. Only the *PDCD1* gRNA had detectable translocations, with two out of three of the translocations being shared between the *S.p.* Cas9 and ABE conditions (Figure 7A). Shared translocations included a fusion of OTE132 to OTE94 and OTE160 to OTE158, with comparable average frequencies ranging between 1.0% - 1.7% and 0.3% - 0.5%, respectively (Figure 7A). The overall estimated translocation burden for this gRNA was estimated to be an average of 1.4% translocations for *S.p.* Cas9 and an average of 2.2% translocations for ABE conditions (Figure 7A). This suggests that translocations are either below 0.1% or not occurring in healthy HSPC donors across higher gRNA specificities. However, it is noteworthy that they are still occurring for both SSB and DSB modalities, as has been previously observed^39^.

**Figure 7.**
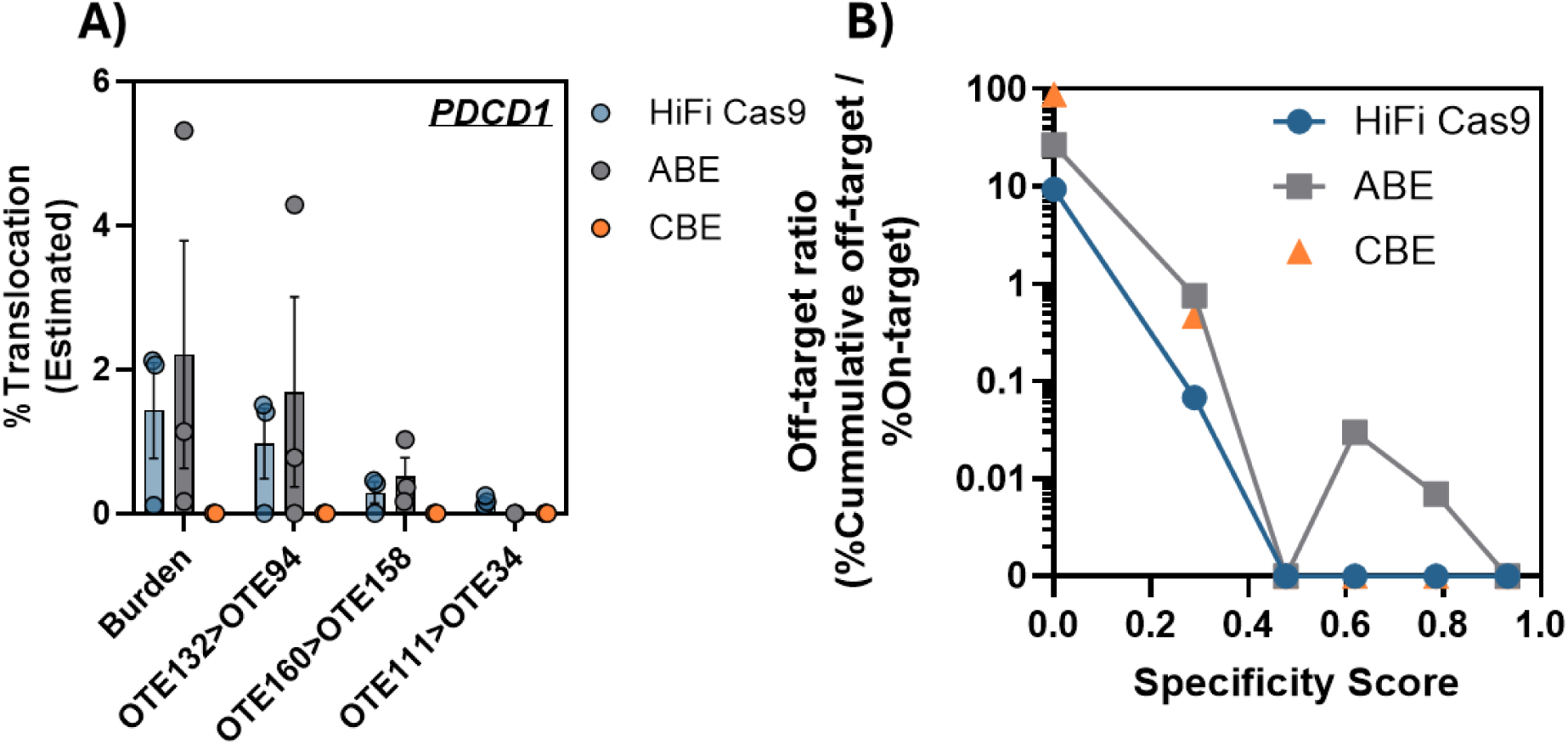
Translocation and cumulative off-target risk. Targeted sequencing data was used for translocation detection between on-target:off-target and off-target:off-target assays in the same pool. **A)** Translocation quantification (FDR < 0.01) at the *PDCD1* site 8 gRNA across different editor modalities in HSPCs and **B)** the cumulative off-target ratio (% cumulative off-target events / % on-target) were calculated for each gRNA in HSPCs with mRNA delivery. Cumulative off-target events for all modalities consider base editing, indel editing, and translocations.

When calculating the normalized risk of cumulative off-target frequencies (indels, base edits, and translocations) across editor modalities throughout our spectrum of gRNA specificities, off-target ratios were observed for the *PDCD1* gRNA over a range 9.4 – 89.0 off-target events per 1 on-target event (Figure 7B). Off-target ratios for the *CYP2C18* gRNA ranged from 0.07 – 0.76 off-target events per 1 on-target event (Figure 7B). Trends consistently showed that overall off-target burden of DSB editors was actually decreased in comparison to SSB base editors for the cumulative frequency of event types monitored using this strategy (Figure 7B). Even though the *B2M* gRNA was considered higher-specificity, a single significant ABE off-target was observed for this treatment contributing to a higher ratio (Figure 7B). This highlights that even higher-specificity gRNAs may generate observable off-targets in clinically relevant cell types.

## Discussion

Our study presents an improved, end-to-end characterized *in cellulo* method for the nomination of off-target sites in CRISPR experiments that we collectively refer to as UNCOVERseq (v1.0). This method leverages several technological improvements to collectively streamline the *in cellulo* nomination process, improve NGS data quality, and increase the number of high-confidence nominated sites through computational analysis improvements compared to other previously published methods. By demonstrating recommended operational conditions that can allow the experiments to be performed independent of cell context with controls grounded in empirical data, we provide a framework to ensure translation to different treatment modalities with quantifiable levels of performance from experiment to experiment. Furthermore, we demonstrate the workflow is capable of nominating relevant unique and shared off-targets for both DSB-based and SSB-based CRISPR editing systems and demonstrate correlations between DSB formation and the frequency of a site to be edited by ABE/CBE editors.

To ensure all relevant off-targets are assessed, high analytical sensitivity is a critical off-target nomination metric. However, accurate calculations of false-negative rates from nomination methods have been challenged by technical difficulties in obtaining an empirically defined gold-standard of all true-positive off-targets. Previous work has led to a perception that *in vitro* biochemical methods are inherently more sensitive than *in cellulo* methods as evidenced by: 1) true positive sites captured by *in vitro* methods like CHANGE-seq that are missed with GUIDE-seq and 2) multiple accounts of *in cellulo* methods being largely a subset of *in vitro* results^6,7^. To reduce risk of false negatives, we demonstrate a strategy for *in cellulo* off-target nomination using UNCOVERseq where high gDNA input and promiscuous editing conditions are used to greatly amplify nomination signal to reproducibly detect sub-0.05% editing events while still retaining sites derived from higher fidelity modalities and primary cell lines. Using UNCOVERseq, we demonstrate that previously published accounts of *in cellulo* methods were not as sensitive as UNCOVERseq. However, it is not clear whether this is due to insufficient operational conditions (read depth, library complexity, etc.) to maximize capabilities of many assays as opposed to the technical improvements that confer enhanced nomination capabilities to UNCOVERseq. We also find that *in vitro* biochemical assays do not sensitively cover the full range of putative true positive lists generated from highly reproduced UNCOVERseq nomination. This provides evidence counter to claims that *in vitro* methods are inherently more sensitive for off-target nomination. Future work should look to further expand gRNAs nominated/confirmed and provide empirical knowledge on the optimal operating conditions for different assays.

Precision is another important metric for off-target nomination methods to appropriately select sites for downstream confirmation. Some methods for off-target nomination can lead to thousands of sites being nominated with low precision which is cost-prohibitive for downstream interrogation given the high coverage depth required for sensitive off-target confirmation. Using UNCOVERseq with sufficient replication (≥3 replicates), we demonstrate that true positives can be obtained from nomination that may approach 100% precision, enabling rapid identification of sites for benchmarking methods. We additionally demonstrate that UNCOVERseq nomination frequencies strongly correlate to observed editing frequencies, and that biochemical *in vitro* methods generally lack as strong of a correlation as compared to *in cellulo* and *in situ* methods. This is an important feature enhancing method-specific site prioritization, as strong correlations to observable frequencies enable better prioritization or filtering of large site lists. Future work should look to better confirm this logic by performing targeted sequencing on more confirmable sites with different levels of reproducibility.

Using a simple prioritization method based on frequency, replication, and high-level indicators of risk (exonic regions vs. intergenic, etc.), we demonstrate UNCOVERseq nominations with recommended experimental structures can result in manageable confirmation panel sizes for downstream confirmation (e.g., <250 sites across a range of specificities). We demonstrate that replicates are important in the off-target nomination process to ensure that sites meeting basic prioritization criteria are appropriately identified, with >25% of putative targets being missed without use of biological replicates. However, to better understand risk after off-target nomination and confirmation across methods, a more standardized scoring system to prioritize off-targets is needed in the future. The fields of oncology and heritable diseases have encountered similar issues and derived guidelines including tiered scoring systems from the American College of Medical Genetics (ACMG) and Association of Molecular Pathology (AMP) and modifications leveraging these criteria^21,40,41^. Gene editing may be able to leverage some of these learnings, but will face unique challenges in categorization of off-target risk since even off-targets in intergenic space during the nomination phase can be at risk for known structural variations derived from DSBs and SSBs. This includes events such as translocations^42^, loss of heterozygosity^43^ (LoH), aneuploidy^44^, and other large variants like multi-kilobase deletions^45^. Given this, it seems likely that probability, frequency, and even proximity to other coding regions will be important criteria for triaging off-targets for assessment. In agreement with previous findings, we find that *in cellulo* methods provide a much stronger relationship between frequency of off-targets nominated and observed compared to *in vitro* methods. This highlights that *in cellulo* methods like UNCOVERseq may have additional utility in future risk scoring criteria given their ability to be predictive of observable frequencies

Benchmarked nomination and confirmation methodologies are needed for both DSB and SSB-based editing modalities. By selecting a variable range of gRNA specificities, we demonstrate that even in popular *ex vivo* models like HSPCs with mRNA delivery, high specificity gRNAs are still sensitive to both SSB indels and base editing off-target effects at frequencies >0.01% and 0.5%, respectively. Furthermore, we demonstrate that indel editing and base editing are rank-order correlated across 34 base editing on/off-targets, supporting the idea that DSB-based nomination methods are effective tools for nominating both indel and base editing activity. Base editing-specific nomination methods, such as CHANGE-seq-BE, have been developed to target base editing events, while demonstrating unique off-target confirmation findings^46^. It is our belief that orthogonal methods for nominating both base editing events and indel events are likely needed for the foreseeable future, especially given some of our findings that some UNCOVERseq-nominated sites generate confirmable indels only in conditions using the SSB base editing modalities in translational cellular contexts.

We envision UNCOVERseq coupled with promiscuous conditions being a powerful tool to help sensitively identify CRISPR-Cas off-targets for interrogation during pre-clinical development phases. By enhancing the quality of off-target nomination and grounding NGS operational conditions in empirical data, we believe that UNCOVERseq improves informed risk assessment of gene editing in translational systems. Furthermore, by detailing inner-workings and described process controls for UNCOVERseq, we hope to enable more flexibility and standardization for the necessary technologies required to operate off-target safety assessment workflows in both research and commercial spaces.

## Methods

### Human cell culture and transfection (K562 and HEK293-Cas9)

K562 (ATCC) and HEK293-Cas9 (ATCC) cells were cultured in Iscove’s Modified Dulbecco’s Medium (IMDM; ATCC) and Eagle’s Minimum Essential Medium (EMEM; ATCC) supplemented with 10% FBS at 37°C with 5% CO_2_. Ribonucleoprotein (RNP) complexes were formed by mixing Alt-R^TM^ *S.p.* Cas9 Nuclease V3 (IDT) or SpyFi^TM^ Cas9 Nuclease (Aldevron) and Alt-R CRISPR-Cas9 sgRNA (gRNA, IDT), and incubating for 20 minutes at room temperature (Molar Ratio: 1:1.2, Cas9:gRNA). For each transfection, 8.0 x 10^5^ cells were washed with 1X phosphate-buffered saline, resuspended in 20 µL of solution SF (Lonza). For K562 cells, RNP complexes at 4 µM were combined with 4 µM of the dsODN (Supplementary Table 1) into the SF solution, while for the HEK293-Cas9 cells, 5 µM gRNA and 0.5 µM dsODN were added to the SF solution. This mixture was transferred into 1 well of a 96-well Nucleocuvette plate (Lonza) and electroporated using program FF-120 (K562) or DS-150 (HEK293-Cas9). Two nucleofections per replicate were performed and each treatment done in triplicate. Following electroporation, cells were transferred to a 6-well plate preheated with either IMDM or EMEM and were incubated at 37°C with 5% CO_2_ for 72 hours. After incubation, genomic DNA (gDNA) was extracted using either the Purelink^TM^ Pro 96 Genomic DNA Purification kit or the Monarch^TM^ Spin gDNA Extraction Kit (New England Biolabs) according to the manufacturer’s instructions, eluted in low-EDTA TE buffer (IDT, 11-05-01-05), and quantified using a NanoDrop 8000 UV-Vis Spectrophotometer (ND-8000-GL).

### Primary T-cell culture and transfection

Frozen human primary pan-T cells (STEMCELL Technologies) from 2 unique human donors were thawed in ImmunoCult-XF T Cell Expansion Medium including 300IU IL-2 (Cytiva) and activated with 10 μL/mL TransAct, human, T cell activator (Miltenyi Biotec) for 48 hours. To prepare for transfection using Lonza 96-well plate 4-D Nucleofector system, cells were counted, pelleted using centrifugation (300 x g, 10 minutes at room temperature), and washed gently with 10 mL 1X phosphate-buffered saline. Cells were again pelleted and resuspended in Lonza Nucleofection Solution P3 at 2.5 x 10^6^ cells/mL. For each electroporation, 5 μL of RNP complex and 3 μL dsODN was added to 20 µL of cells in P3 (5 x 10^5^ cells/nucleofection) for a final concentration of 4 μM RNP (1:1.2 ratio of Cas9 to gRNA) and 1-4 μM dsODN. Where tag was not included, 3 μL of IDT Alt-R Cas9 Electroporation Enhancer was added for 3 µM final concentration to achieve a fixed final nucleofection reaction volume of 28 μL. Each reaction was mixed by pipetting and 25 µL was transferred to an electroporation cuvette plate. The cells were electroporated according to the manufacturer’s protocol using the Amaxa 96-well Shuttle and nucleofection protocol 96-EH-140. After electroporation, the cells were resuspended in 75 μL pre-warmed IL-2 culture media in the electroporation cuvette. Triplicate aliquots of 25 μL of recovered cells were further cultured in 175 μL pre-warmed IL-2 media with TransAct. Cells were incubated for 72 hours, after which gDNA was isolated and quantified.

### iPSC culture and transfection

iPSCs from fibroblasts (Coriell Institute, GM23338) were cultured in mTeSR™ Plus media (Stemcell Technologies) at 37°C with 5% CO_2_. RNPs were formed as described above. For transfection using Lonza 96-well plate 4-D Nucleofector system, cells were detached using ReLeSR™ (Stemcell Technologies) and washed with 1X phosphate-buffered saline. Cells were resuspended in P3 buffer at 2 x 10^5^ cells/nucleofection. CRISPR reagents at required final concentrations (4 µM RNP; 0.5 µM dsODN) were added to the mix to make a final volume of 25 µL, and of which 20 µL was transferred to the nucleocuvette for electroporation. The nucleovette plate was electroporated using code CA-137. After the nucleofection, cells were recovered and plated in complete mTeSR Plus medium with 1X CloneR™ 2 supplement (Stemcell Technologies). Recovery media was added to the electroporated cells to achieve a final volume of 100 µL, and 25 µL of this was added to 175 µL media per replicate well for final plating in a vitronectin-coated 96-well plate. During recovery and growth at 37°C with 5% CO_2_ for up to 96 to 120 hours, media changes were performed as desired and/or following manufacturer’s protocols for media and CloneR 2 supplement. gDNA extraction and quantification was performed as described above.

### Off-target Nomination with UNCOVERseq

500 ng of purified gDNA was enzymatically fragmented and adapter-ligated using the xGen^TM^ DNA Library Prep EZ UNI kit along with the xGen Deceleration Module (IDT, xGen DNA Library Prep EZ UNI 96 rxn, 10009822; xGen Deceleration Module 96 rxn, 10009823) according to the manufacturer’s instructions and cleaned with AMPure XP beads (Beckman). Following fragmentation and adapter ligation, rhPCR was performed using rhAmpSeq^TM^ Library Mix 1 (IDT) to amplify the DNA in a single tube using a forward primer specific to the P5 adapter, a reverse primer specific for top and bottom strand of the integrated dsODN tag, and an adapter-blocking oligo corresponding to each strand of the dsODN^31^. Following PCR, samples were diluted 1:40 with nuclease-free water and used in a second PCR with rhAmpSeq^TM^ Library Mix 2 (IDT) that added a unique P7 adapter to each library. Libraries were then cleaned with AMPure XP beads and run on an Agilent Fragment Analyzer for library quality assessment. All libraries were quantified with the Qubit 1X dsDNA HS Assay kit (Invitrogen) and pooled in equimolar amounts. All libraries were sequenced using an Illumina MiSeq or NextSeq2000 instrument with 150-bp paired-end reads. All oligonucleotides used in preparation of UNCOVERseq libraries are listed in Supplementary Table 1.

### Vector Construct and *in vitro* transcription of modified mRNA

Genes encoding HiFi Cas9^47^, ABE8e^37^, AncBE4max^36^, and PE2^28^ were each cloned into a plasmid vector with a dT7 promotor followed by a 5’UTR, Kozak sequence, ORF, and 3’UTR for subsequent in vitro transcription. Chemically modified Cas9, ABE8e, AncBE4max and PE2 mRNA was transcribed *in vitro* from the PCR templates with full substitution of uridine by N^1^-methylpseudouridine. Co-transcriptional capping was achieved using CleanCap AG analog (TriLink), yielding a 5′ Cap 1 structure. Transcription reactions were carried out with the HiScribe T7 High Yield RNA Synthesis Kit (New England Biolabs) in 0.5× transcription buffer supplemented with 4 mM CleanCap AG. Following synthesis, mRNAs were purified using the Monarch Spin RNA Cleanup Kit (New England Biolabs). PCR templates incorporated mammalian-optimized UTRs (TriLink) and a 120-nucleotide poly(A) tail.

### HSPC culture and transfection

CD34⁺ HSPCs from a single human donor (Fred Hutch Cancer Center, RO04089) were cultured at 1 × 10⁵ cells/mL in StemSpan SFEM II medium supplemented with 100 ng/mL SCF, 100 ng/mL TPO, 100 ng/mL FLT3L, 100 ng/mL IL-6, 20 µg/mL streptomycin, and 20 U/mL penicillin. Cultures were maintained at 37 °C in a humidified incubator with 5% CO₂. For electroporation, Cas9 mRNA and gRNA were mixed at a 1:1 weight ratio (3 µg each per reaction). ABE, CBE, and PE2 mRNAs were used at equimolar amounts to Cas9 (3.4 µg ABE, 4 µg CBE, and 6 µg PE2), and pegRNA was added at the same molar amount as gRNA (4.2 µg). HSPCs were resuspended in 20 µL Lonza P3 buffer and electroporated using a Lonza 4D-Nucleofector (program DZ-100). After electroporation, cells were plated at 1 × 10⁵ cells/mL in HSPC medium.

### Computational Analysis – Nomination

Following NGS, Illumina adapters and UMIs were identified and annotated using Picard MarkIlluminaAdapters. Tag sequences were identified and trimmed using Cutadapt v4.2^48^. Sequencing reads were aligned to hg38 (GRCh38.p12) reference genome using BWA mem^49^ v0.7.15 and UMI consensus reads were generated based on consensus from a single-strand (minimum UMI consensus size = 1) using fgbio v0.7.0 (https://github.com/fulcrumgenomics/fgbio). Nomination of candidate off-target sites began by using mapped UMI consensus reads to create a flanked search space (+/- 40 bp) to perform alignment between the guide and empirical target region using a glocal implementation of the Needleman-Wunsch alignment^50,51^. After a candidate match to the gRNA spacer region was identified in the sequencing data, nominated off-target sites were identified using a hypergeometric test with multiple testing correction (Benjamini & Hochberg; FDR<0.05) by comparing individual treatment samples and pooled control samples for significant differences in representation between the two. We used the following criteria to nominate off-target sites from this analysis for verification: 1) at least one sample nominated a given site with NGS evidence on both sides of the cut site, 2) Levenshtein distance < 7 as determined post-alignment, and 3) significant adjusted p-value when comparing the frequency of the event to the pooled control(s). Nominated on/off-target sites had additional meta-data added based on alignment/genomic context and were placed into described Tiers based on this meta-data (Supplementary Table 2).

### Library Preparation – Confirmation

Genomic DNA was extracted from control and genome-edited cells as described above. Libraries for amplicon NGS were prepared using a previously described rhAmpSeq amplification-based method (IDT) using 100 ng of gDNA input^52^. Briefly, the first round of PCR was performed using target-specific primers. A second round of PCR was used to incorporate P5 and P7 Illumina adapters to the ends of the amplicons for universal amplification. Libraries were purified using Agencourt AMPure XP system (Beckman Coulter, Brea, CA, USA) and quantified by quantitative PCR (qPCR) before sequencing on the Illumina MiSeq platform (v.2 chemistry, 150-bp paired end reads; Illumina). Read demultiplexing was performed on the resulting BCL files using Picard v2.18.9 (https://github.com/broadinstitute/picard) IlluminaBasecallsToFastq.

### Computational Analysis – Confirmation

Analysis of the sequencing data to identify confirmed off-target editing at the nominated sites was performed using CRISPAltRations v1.2.1^53^. This analysis comprised two parallel workflows: identification of indels at the position of the DSB/SSB, and identification of base editor-induced A->G (ABE) or C->T (CBE) transversions in the relevant base-editing window.

For identifying indels, the window for event quantification was centered on the canonical cut site and events quantified utilizing the default window size for Cas9 (8 bp). To determine whether indels found in the sequencing data could result from bona fide off-target cleavage, indels were grouped by location relative to the cut site (prioritizing minimum distance to cut site) followed by fitting counts of events to a negative binominal model with a Wald test for significance in each location bin per off-target using the DESeq2 package^35^ within IDT’s OTEasy tool (Schmaljohn et al., Manuscript in Preparation). For classification of indel off-target editing, the tool requires: 1) sufficient read coverage for the site (>1,000x) in all replicates, 2) significant edits to occur at or adjacent to the cut site after optimal alignment, 3) the classified cumulative significant edits to exceed 0.01%, 4) the comparison of treatment/control samples at the site to have a significant adjusted p-value (p < 0.05), and 5) an average coverage frequency of at least 5x the ascribed cumulative frequency observed (e.g., for 0.1% editing, at least 5,000x coverage). Blunt dsDNA tag integration quantification for quality control analysis were classified by aligning quantified insertions >10 bp in size from the CRISPAltRations output to the expected dsDNA tag using the biopython implementation of the Needleman-Wunsch aligner^50^. Alignments with an alignment score greater than 50 were quantified as a tag integration event, and events were considered imperfect if any base was mutated or missing from the expected dsDNA sequence alignment.

For identifying base editing-generated off-target effects, the window for event quantification was centered in the middle of canonical base-editing window between position +5/+6 of the spacer (5’ to 3’) with a 5 bp window for quantification. To determine significant base-editing transitions resulting in off-target editing, all individual events that contained an ABE (A>G or T>C) or CBE (C>T or G>A) transition were grouped according to unique base editing events in the window and fitting counts of events to a negative binominal model with a Wald test for significance in each location bin per off-target using the DESeq2 package^54^ within IDT’s OTEasy tool (Schmaljohn et al., Manuscript in Preparation). For classification of adenine base editing at off-targets, the tool requires: 1) sufficient read coverage for the site (>1,000x) in all replicates, 2) the classified cumulative significant edits to exceed 0.5%, 3) the comparison of treatment/control samples at the site to have a significant adjusted p-value (p < 0.05), and 4) an average coverage frequency of at least 5x the ascribed cumulative frequency observed.

### Computational Analysis – Translocations

To quantify translocations from editing, Primer Anchored Statistical Translocation Analysis (PASTA) was used (Kurgan et al., Manuscript in Preparation). This analysis is only performed on the amplicon sequencing pools containing the on-target edit (because multiplexed amplification is a requirement for event detection using the method), and reactions not containing the on-target edit are unlikely to have any significant translocation events. To quantify translocations, expected primers were identified in reads using fg-idprimer (https://github.com/fulcrumgenomics/fg-idprimer; -k=6, -K=8, -S=5, --max-mismatch-rate=0.07). Following this, treatment/control pairs had their counts paired and primer count frequencies subjected to a one-tailed hypergeometric test with Benjamini-Hochberg correction (statsmodel v0.15.0; default settings) to calculate an adjusted p-value (p-adj). Unexpected primer pairs with padj < 0.01 with no flags were classified as a translocation and had the translocation frequency (P) calculated using the following equation:

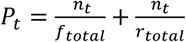

Where ‘n’ is equal to the count of the unexpected primer pair of interest, ‘t’ is the significant translocation being interpreted, ‘f’ is the total count of the shared forward primer events excluding the count participating in the ‘n’ translocation event, and ‘r’ is the total count of shared reverse primer events excluding the count participating in the ‘n’ translocation event. The translocation frequency is then adjusted by the background level frequency in the control by subtracting any translocation frequency observed in the control sample from the treatment frequency. Total translocation burden (B) was calculated using the following equation:

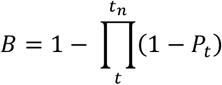

Where ‘t’ is equal to a significant translocation, and ‘tn’ is equal to the last significant translocation of all translocations. All translocations for the purposes of this equation are assumed to be occurring independently. Using the method, translocations are quantified if: 1) the estimated frequency exceeds 0.1% of editing, 2) the translocation has a significant p-value (p < 0.01), and 3) the translocation is found to meet these criteria in all replicates.

### Data Availability Statement

Data generated and analyzed during this study using the UNCOVERseq method are available for download. These datasets are provided exclusively for **non-commercial research purposes**. Use of the data for any commercial applications—including but not limited to training or developing predictive models, designing tools for gene editing applications, or integrating into commercial software or services—is strictly prohibited without prior written permission from the authors. This restriction applies retroactively to all versions of the datasets provided in this work, including those downloaded prior to the publication of this statement. Users who have previously accessed the data are requested to comply with these updated terms and refrain from any commercial use, without prior written permission.

### Disclaimers and Conflicts of Interest

Products and tools supplied in this manuscript by IDT are for research use only and not intended for diagnostic or therapeutic purposes. Purchaser and/or user are solely responsible for all decisions regarding the use of these products and any associated regulatory or legal obligations. For informational use only. The data provided are for informational use only and should not be used as the sole basis for any critical decision making. The data generated are based on assay procedures that have not undergone full validation: formal design and development activities are on-going. K.J.K., B.T., M.L.S., G.L.K., S.C., R.C., E.S., T.O., K.M., A.S.P., H.Z., M.B., M.M., S.W., R.T., G.R., and A.M.J. are employees of IDT, which offers reagents and services for sale similar to some described in the manuscript. M.K.C. has equity in Kamau Therapeutics.

---

**Supplementary Figure 1.**
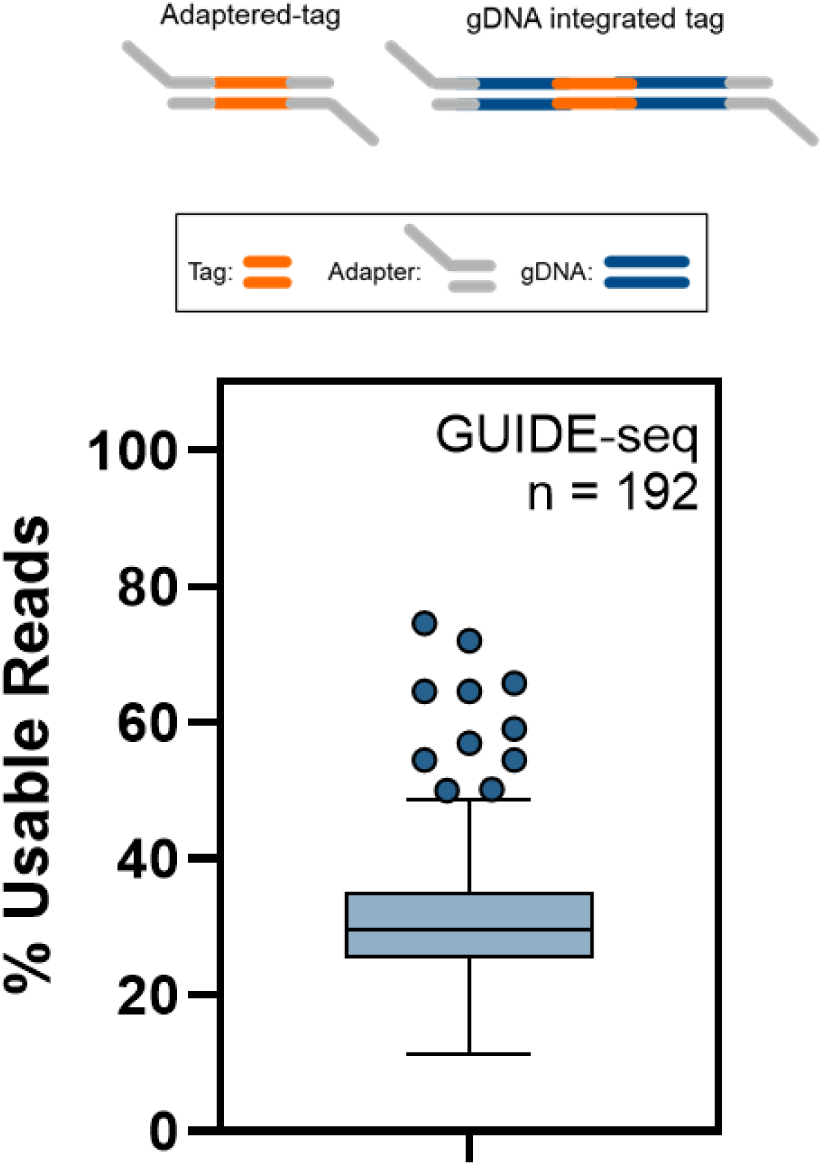
Quantification of read loss due to adaptered tag in the GUIDE-seq protocol. A depiction is shown regarding what kind of events form during library preparation with GUIDE-seq, with the usable reads (non-dsDNA:adapter reads) measured across nominating off-targets for 192 gRNAs in K562 (n = 1 per gRNA).

**Supplementary Figure 2.**
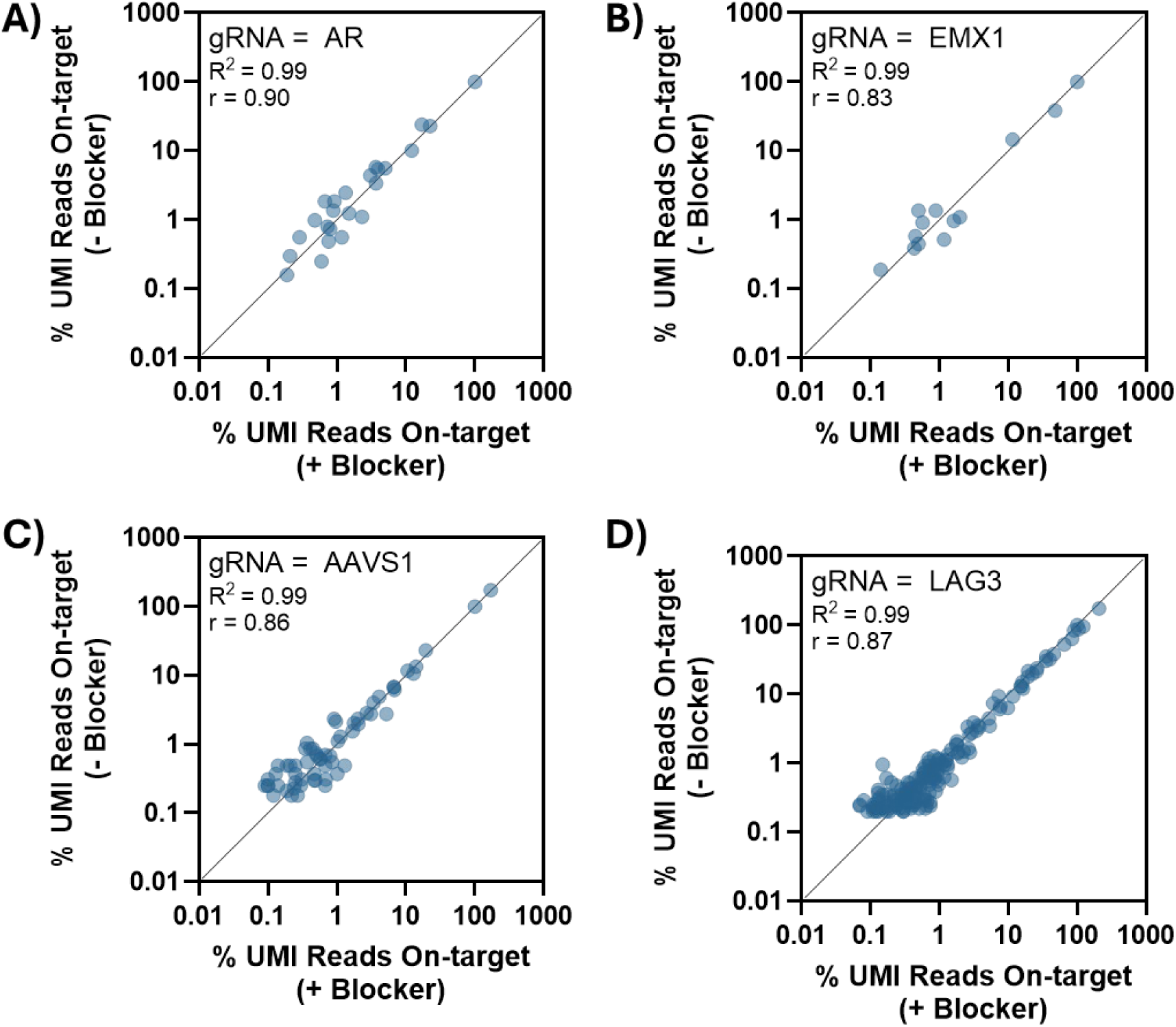
Comparison of UNCOVERseq nomination frequencies with and without dsDNA tag:adapter blocker. Comparison of average nomination frequencies (normalized to the UMI-corrected on-target frequency) in K562 (n = 3 per gRNA) with and without the dsDNA tag:adapter blocker (Blocker) from shared sites for the following gRNAs: **A)** *AR*, **B)** *EMX1*, **C)** AAVS1, and **D)** *LAG3*.

**Supplementary Figure 3.**
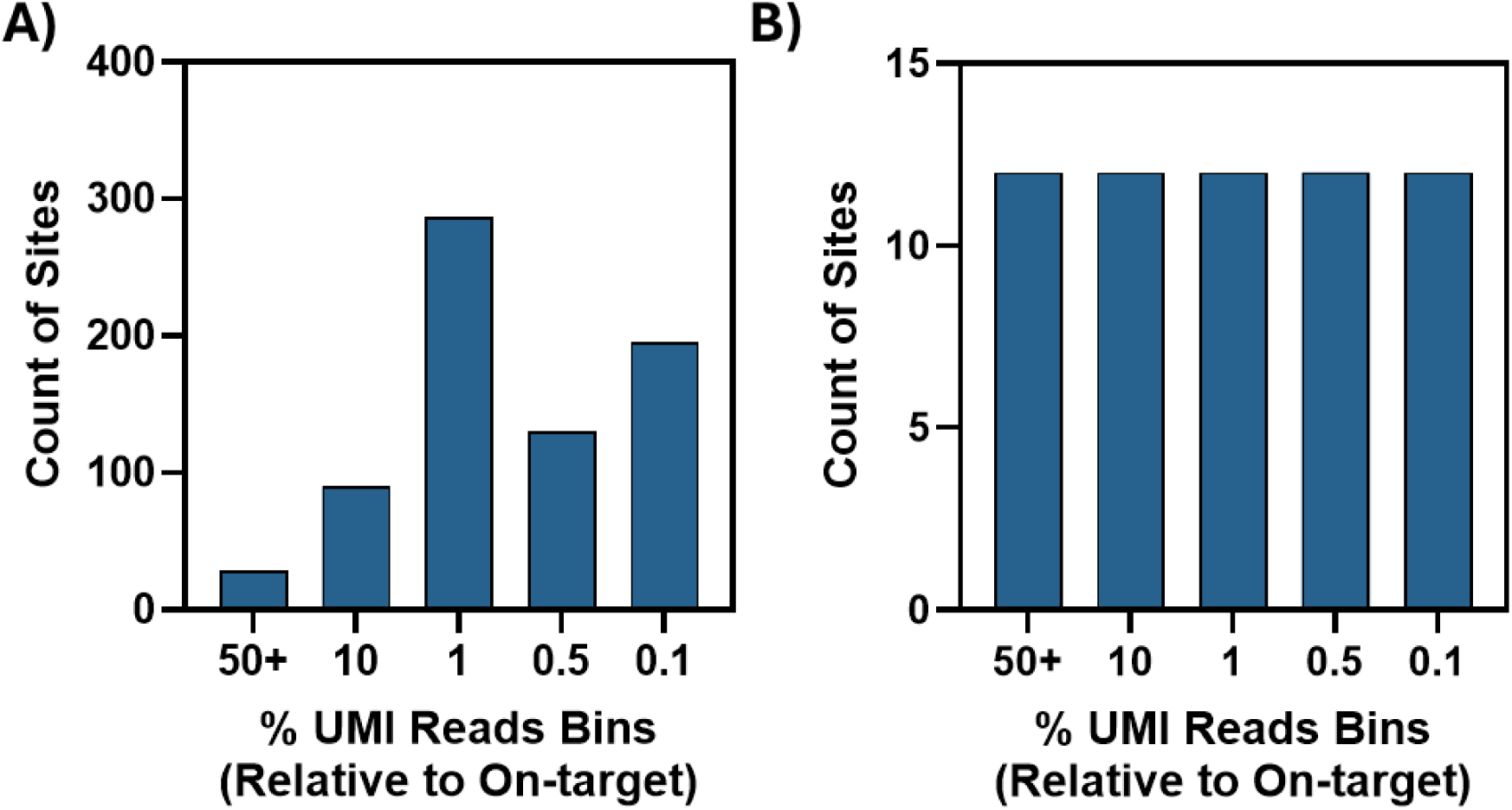
Site selection for LAG3 process control panel. UNCOVERseq was used to nominate targets of the *LAG3* site 9 gRNA in HEK293-Cas9 (n = 12 biological replicates) and **A)** the average nomination frequency (normalized to the on-target frequency) of each site was binned into 5 frequency bins (0.10 – 0.49%; 0.50% – 0.99%; 1 - 9.9%; 10 - 49.9%; >50% UMI reads relative to the on-target). **B)** Selected sites per frequency bin for interrogation as a part of process control 60-plex.

**Supplementary Figure 4.**
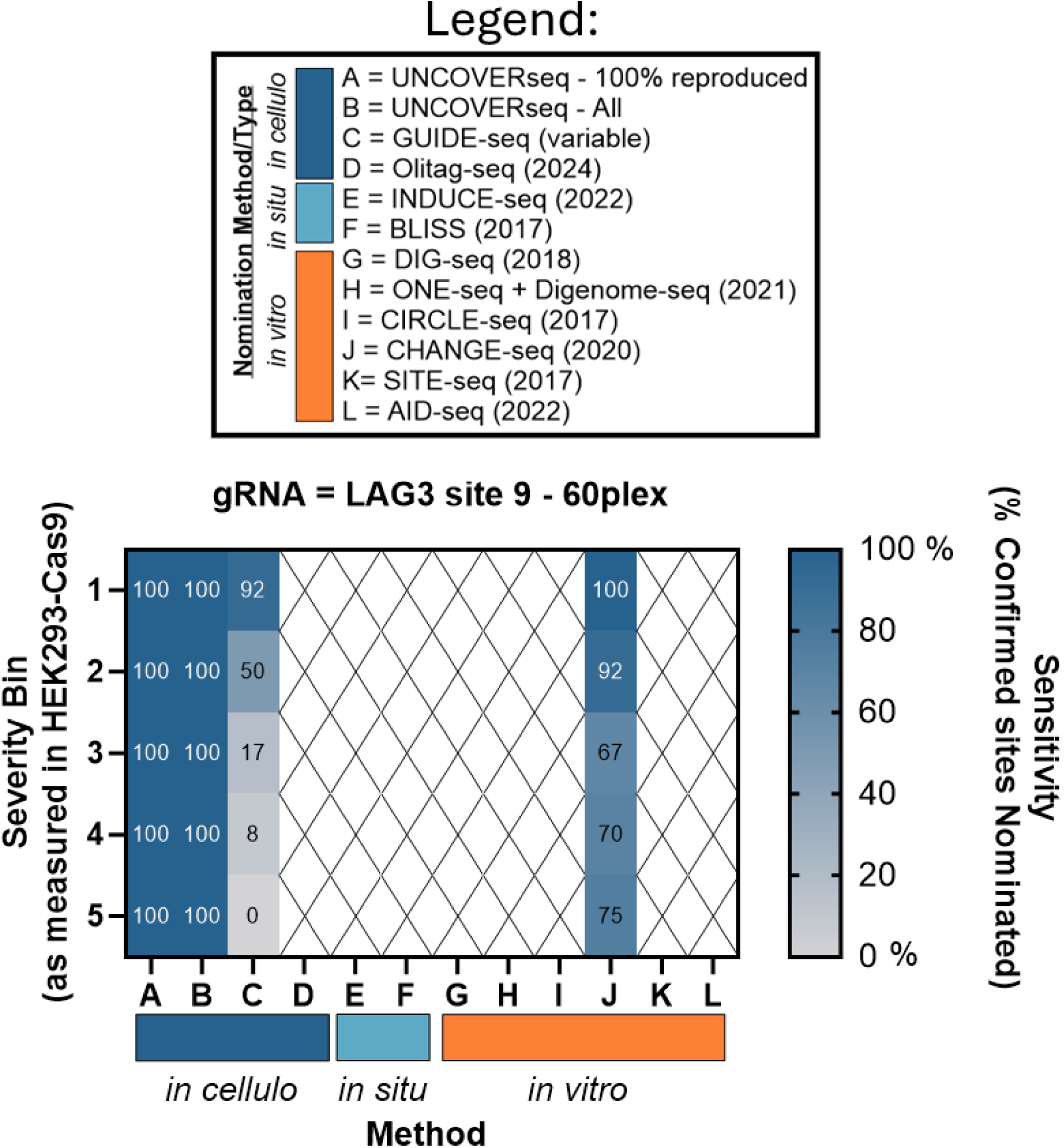
Comparative analysis of UNCOVERseq at the confirmed LAG3 60-plex site list to other nomination technologies. Comparison of the sensitivity of other published accounts of nomination technologies to nominate sites previously confirmed UNCOVERseq off-target sites at the *LAG3* site 9 (n = 12 nomination replicates) gRNA.

**Supplementary Figure 5.**
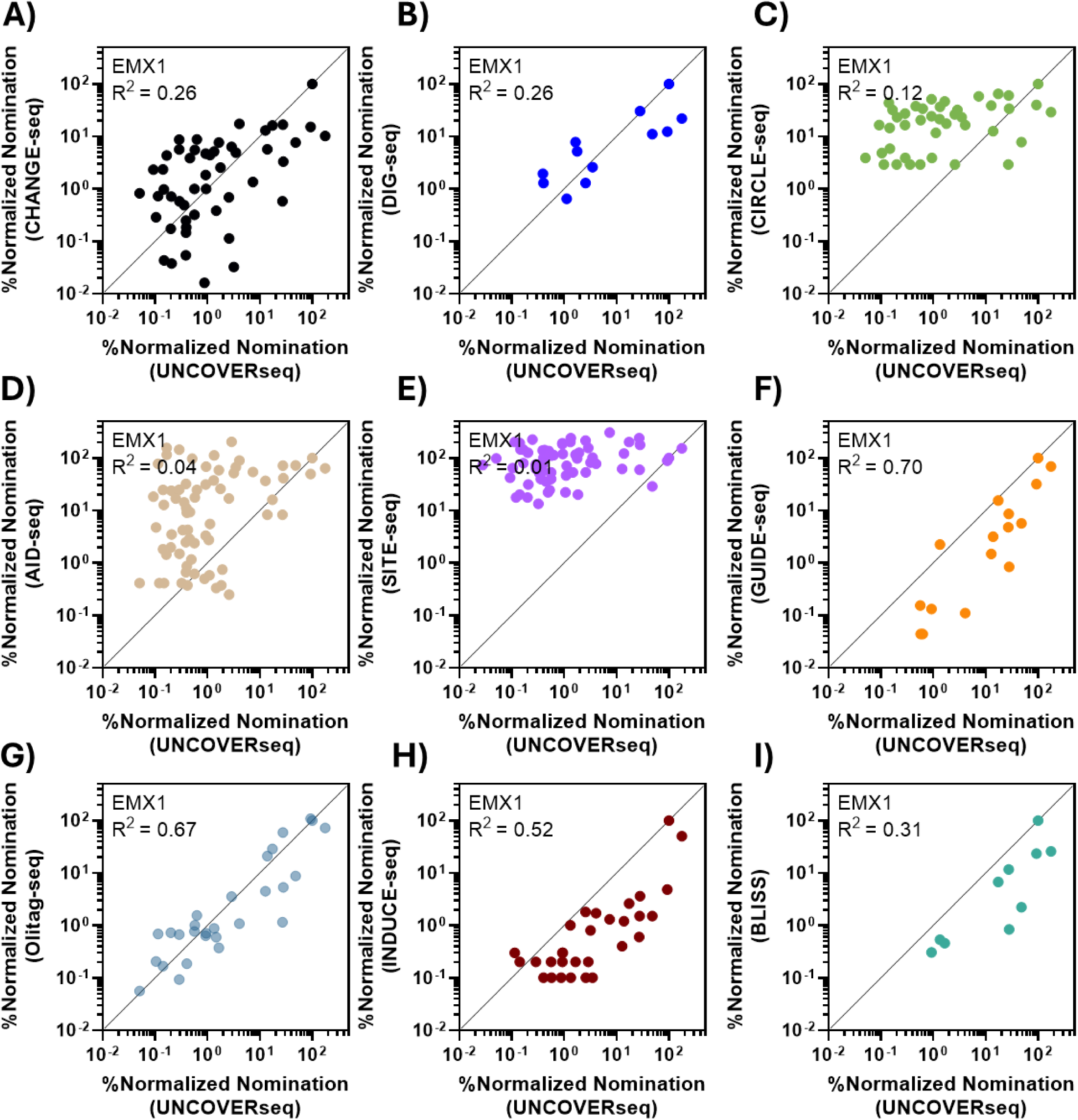
Comparative analysis of UNCOVERseq nomination frequencies to other previously published technologies at the EMX1 site. Comparison of the nomination frequencies from shared sites of published accounts of nomination technologies (normalized to the on-target frequency) to average nomination frequency of fully reproduced EMX1 UNCOVERseq nomination (n = 6 replicates). Biochemical *in vitro* methods displayed are **A)** CHANGE-seq **B)** DIG-seq **C)** CIRCLE-seq **D)** AID-seq and **E)** SITE-seq. Cell-based *in cellulo* or *in situ* methods displayed are **F)** GUIDE-seq **G)** Olitag-seq **H)** INDUCE-seq and **I)** BLISS

**Supplementary Figure 6.**
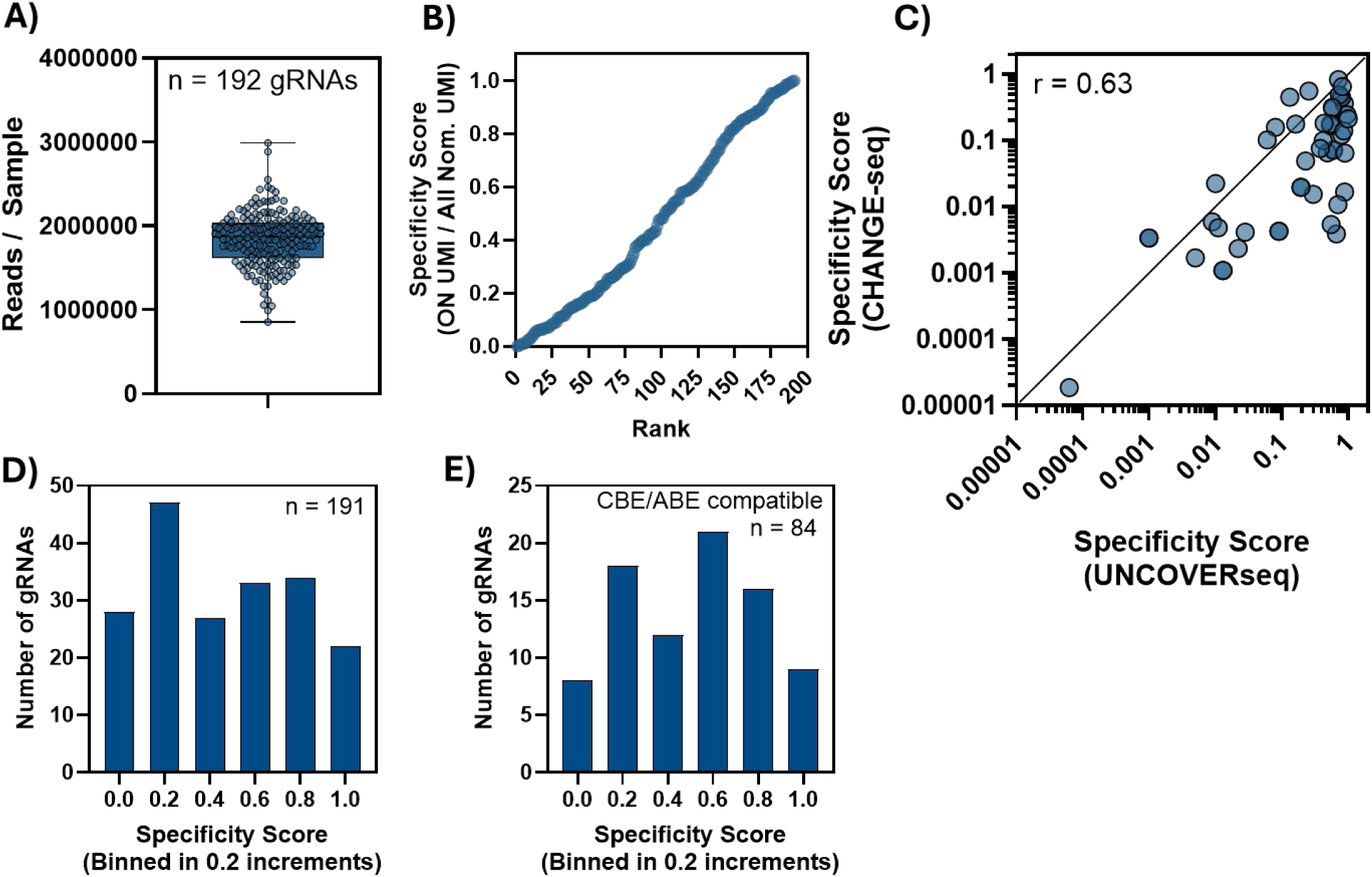
UNCOVERseq gRNA specificity scores for nominated gRNAs and ABE/CBE-compatible gRNAs. 192 gRNAs were individually transfected into HEK293-Cas9 to perform UNCOVERseq. **A)** Read depth per sample of each gRNA and **B)** the rank-order specificity score were quantified. **C)** Comparison of derived specificity scores to those from the published CHANGE-seq method (2020). Specificity scores were binned from 0 to 1 in 0.2 increments and the # of gRNAs per binned counted for **D)** all gRNAs and **E)** gRNAs that met ABE criteria (at least one “A” in 5’ position 4 – 7) and CBE criteria (at least one “C” in 5’ position 4 – 8).

**Supplementary Figure 7.**
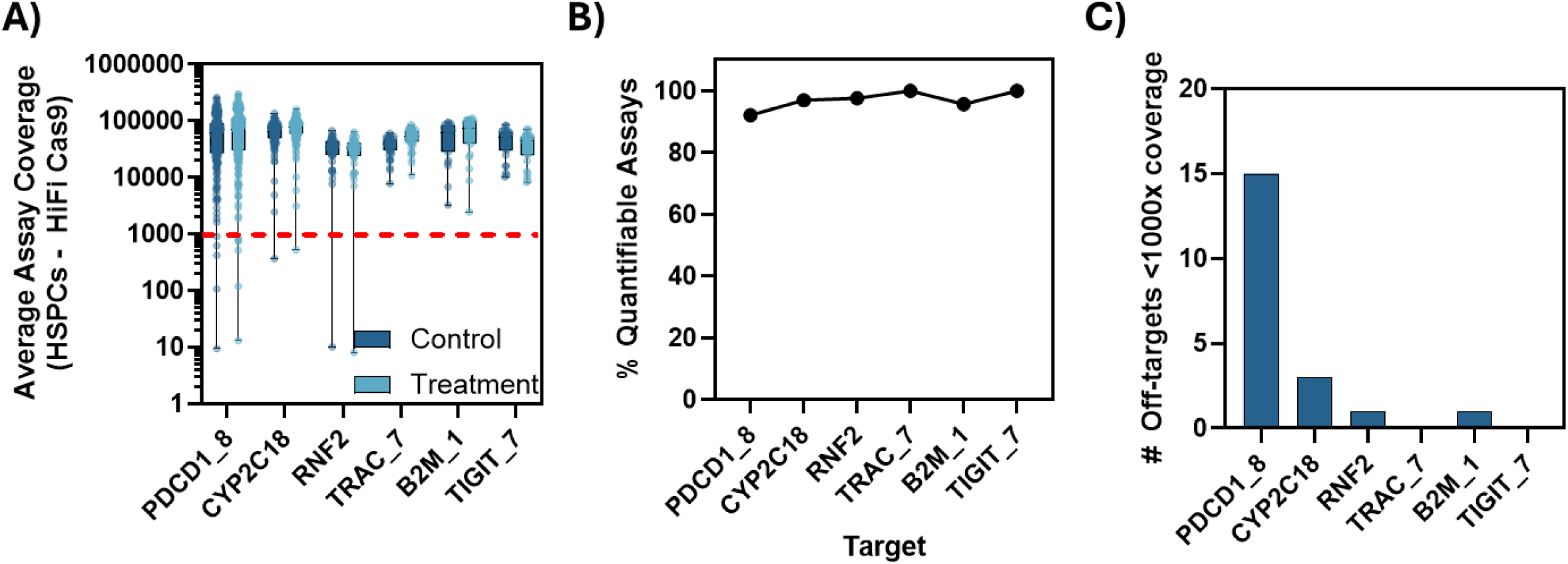
Sequencing performance per gRNA off-target panel. **A)** Average targeted sequencing coverage for each assay within the multiplexed rhAmpSeq panel created for confirmation of off-targets is plotted using paired treatment/controls for the HSPC HiFi Cas9 editing condition (n = 3 per treatment) with the >1,000x coverage requirement depicted (dotted red line). **B)** Frequency of all replicates reaching >1,000x for each target in the panel and **C)** the number of targets failing to meet this threshold is quantified per gRNA.

**Supplementary Figure 8.**
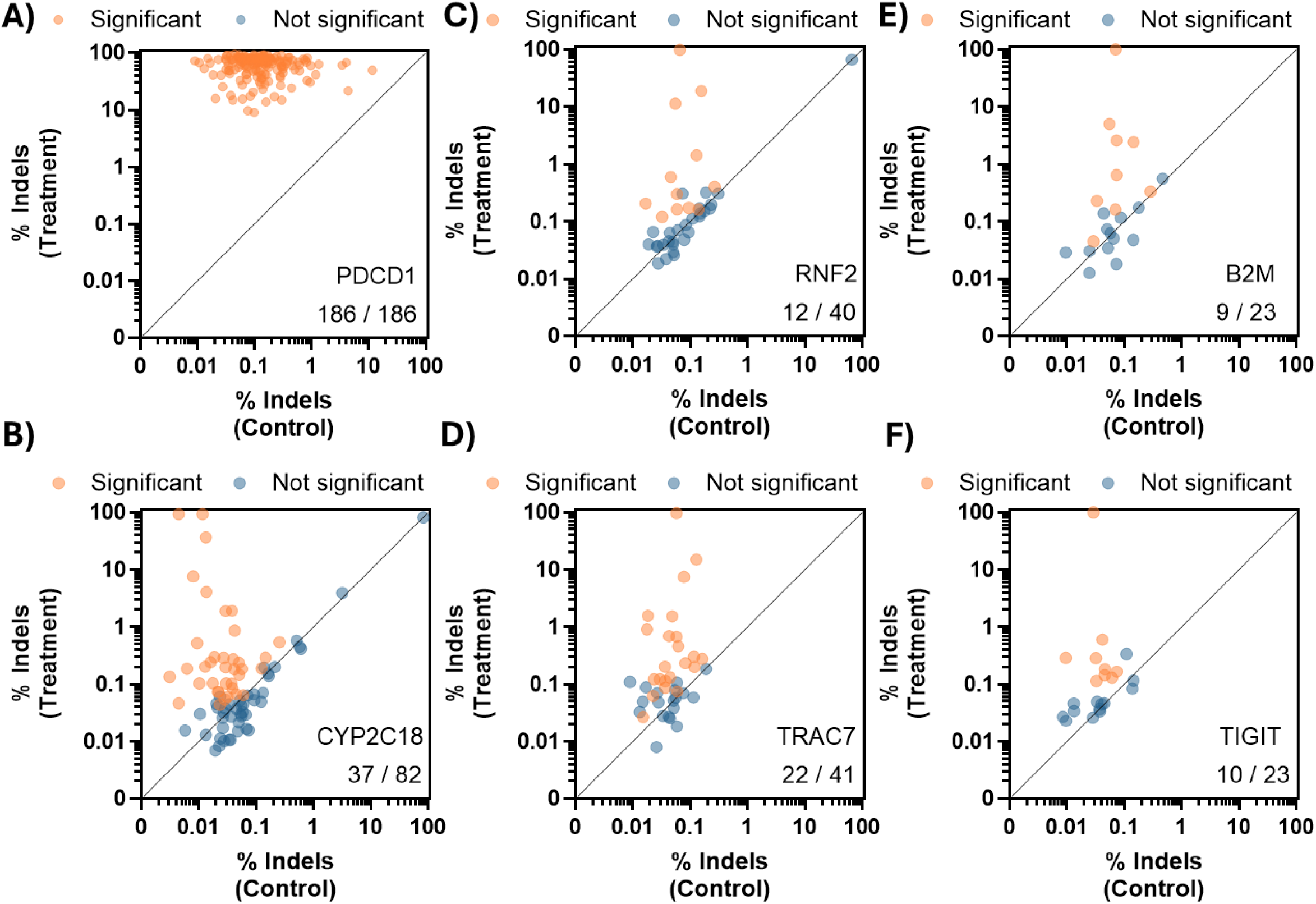
Confirmation in HEK293–Cas9 (indels). Editing quantification at assays with >1,000x coverage. Each dot represents a gRNA on/off-target with the average raw frequency of indels of the control (x-axis) and treatment (y-axis) plotted. Blue dots indicate sites with no statistical significance while orange dots indicate significant sites (p adj<0.05) for the **A)** *PDCD1* site 8, **B)** *CYP2C18*, **C)** *RNF2*, **D)** T*RAC* site 7, **E)** *B2M* site 1, and **F)** *TIGIT* site 7 gRNAs. Text at the bottom right displays the total number of confirmed sites out of all interrogated with sufficient depth (>1,000x).

**Supplementary Figure 9.**
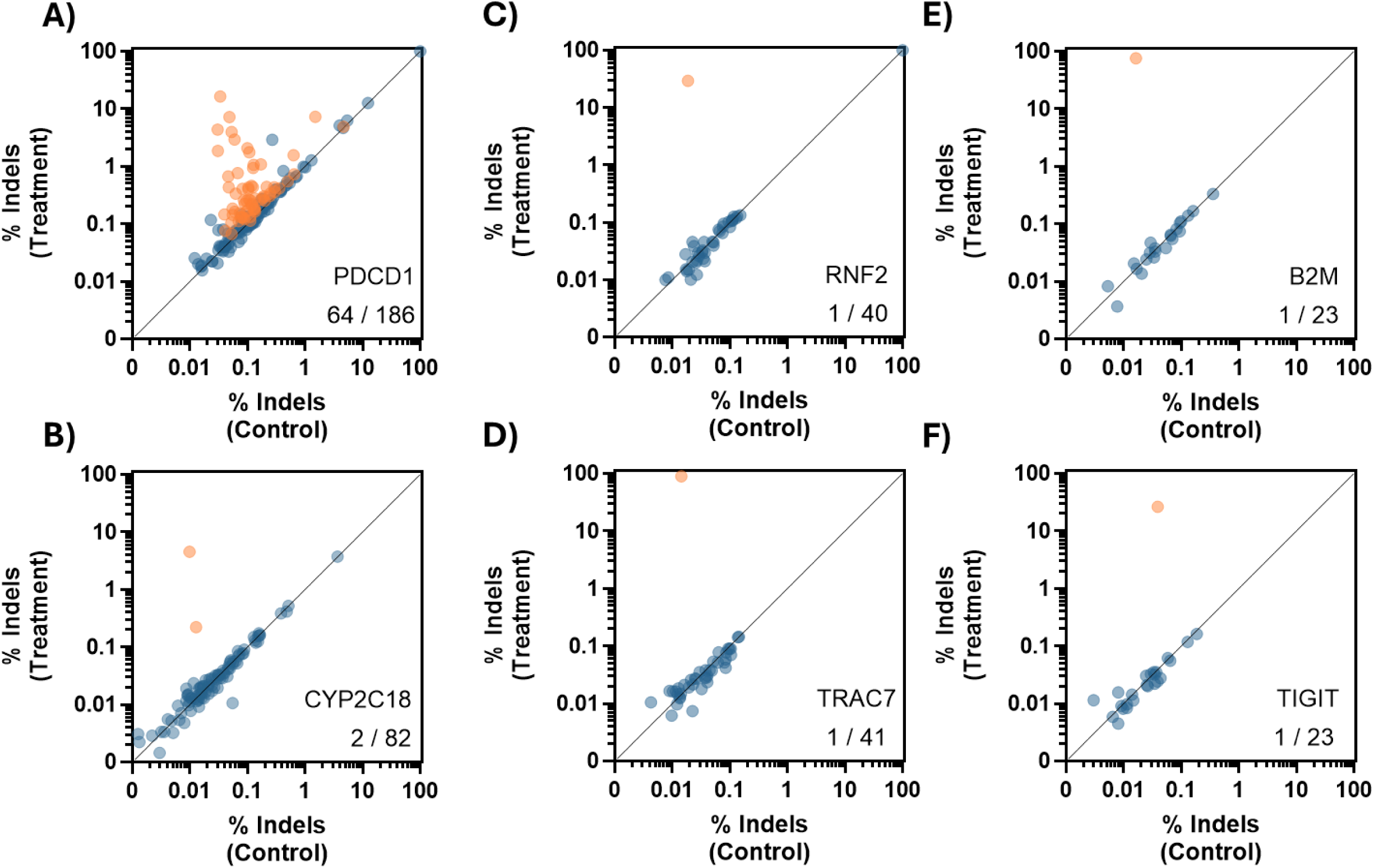
Confirmation in HSPCs with HiFi Cas9 delivered as mRNA (indels). Editing quantification at assays with >1,000x coverage. Each dot represents a gRNA on/off-target with the average raw frequency of indels of the control (x-axis) and treatment (y-axis) plotted. Blue dots indicate sites with no statistical significance while orange dots indicate significant sites (p adj<0.05) for the **A)** *PDCD1* site 8, **B)** *CYP2C18*, **C)** *RNF2*, **D)** T*RAC* site 7, **E)** *B2M* site 1, and **F)** *TIGIT* site 7 gRNAs. Text at the bottom right displays the total number of confirmed sites out of all interrogated with sufficient depth (>1,000x).

**Supplementary Figure 10.**
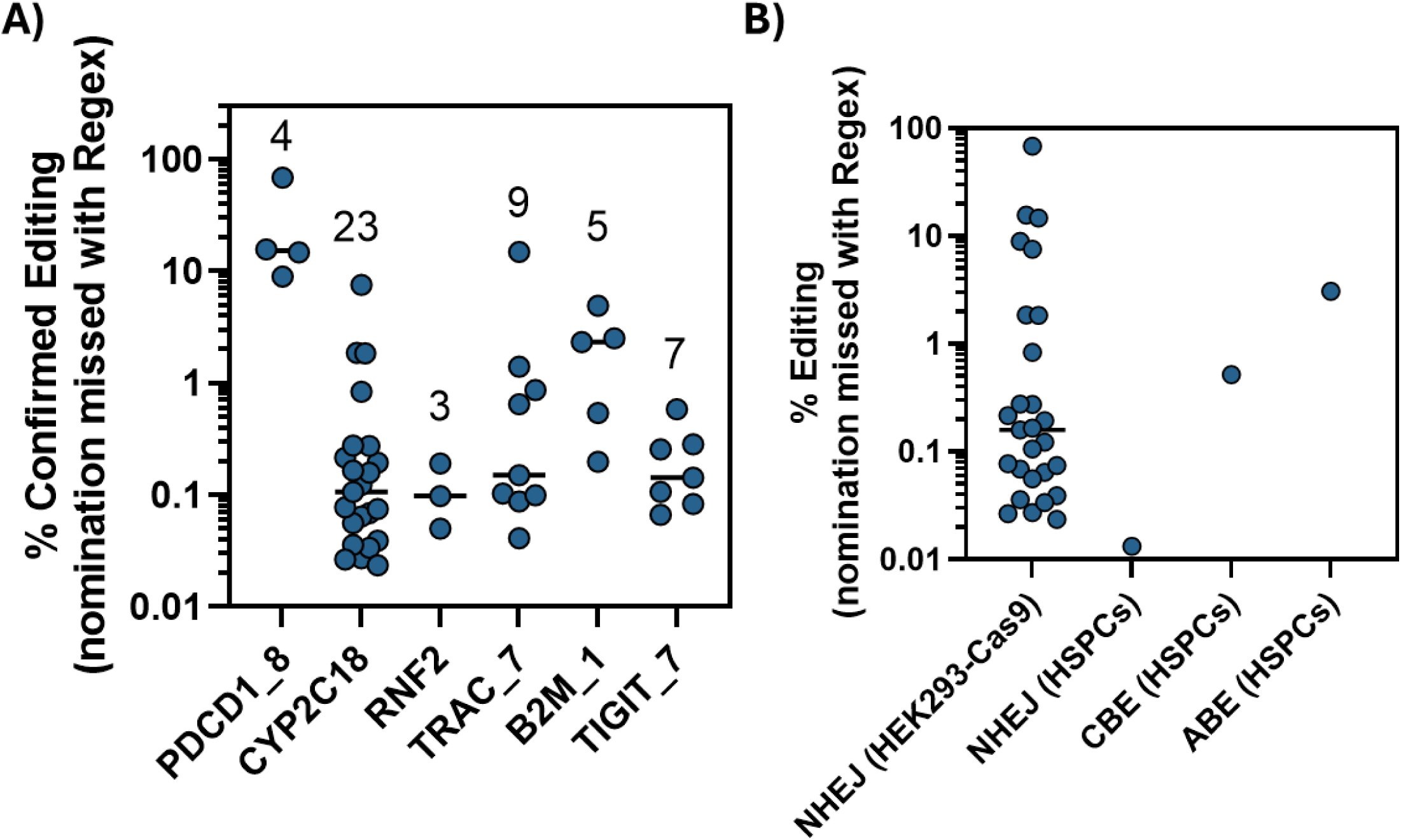
Confirmable editing sites that can be missed with Regex alignment approach. Confirmed off-targets (p-value < 0.05) were intersected with those missed using the Regex method for determining off-target alignment distance < 7 and frequencies of these sites determined **A)** per gRNA in HEK293-Cas9 and **B)** per condition, between HEK293-Cas9 and HSPCs (HiFi Cas9, CBE, and ABE).

**Supplementary Figure 11.**
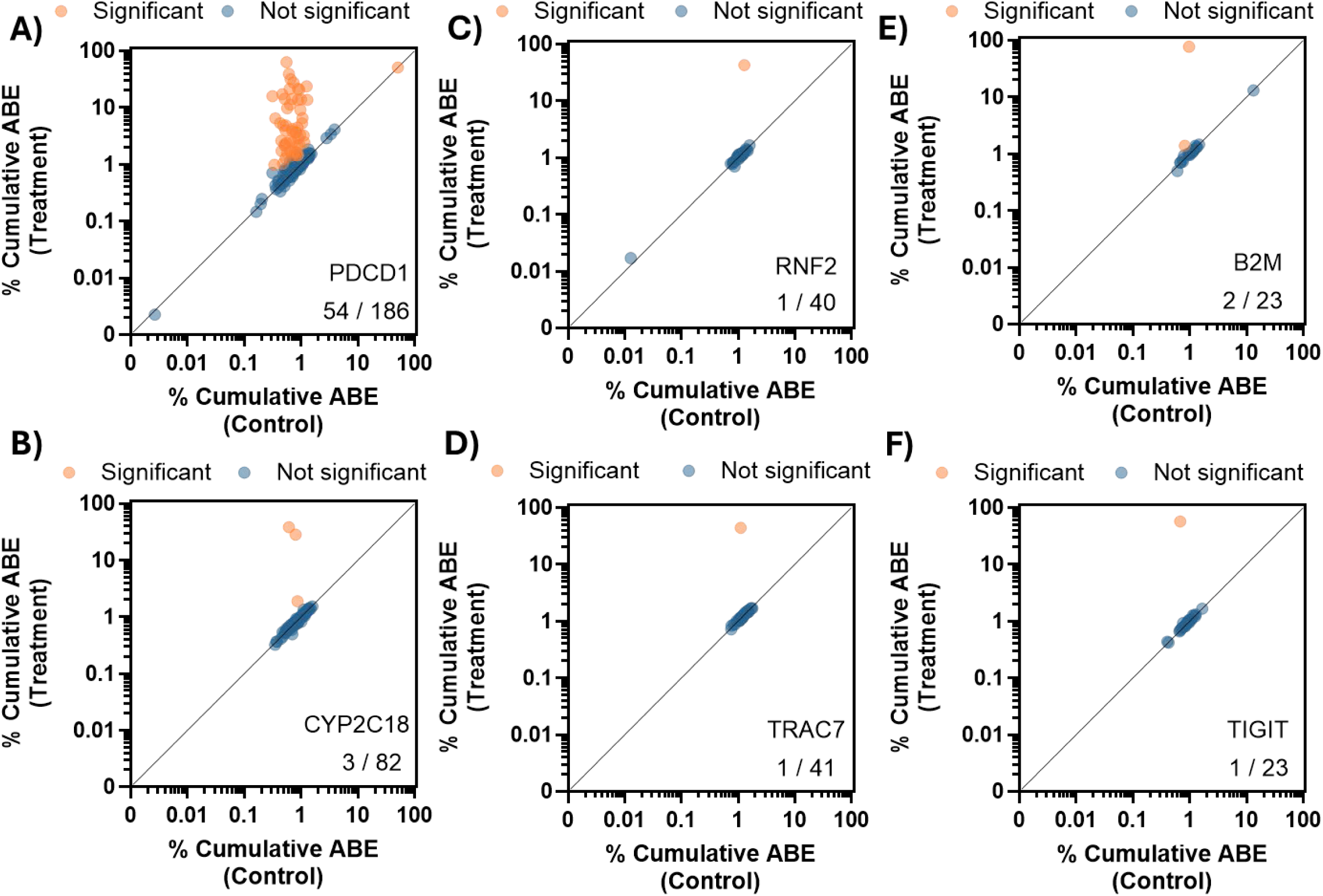
Confirmation in HSPCs with wildtype *S.p.* Cas9-ABE delivered as mRNA (ABE). Editing quantification at assays with >1,000x coverage. Each dot represents a gRNA on/off-target with the average raw frequency of cumulative ABE transition events of the control (x-axis) and treatment (y-axis) plotted. Blue dots indicate sites with no statistical significance while orange dots indicate significant sites (p adj<0.05) for the **A)** *PDCD1* site 8, **B)** *CYP2C18*, **C)** *RNF2*, **D)** T*RAC* site 7, **E)** *B2M* site 1, and **F)** *TIGIT* site 7 gRNAs. Text at the bottom right displays the total number of confirmed sites out of all interrogated with sufficient depth (>1,000x).

**Supplementary Figure 12.**
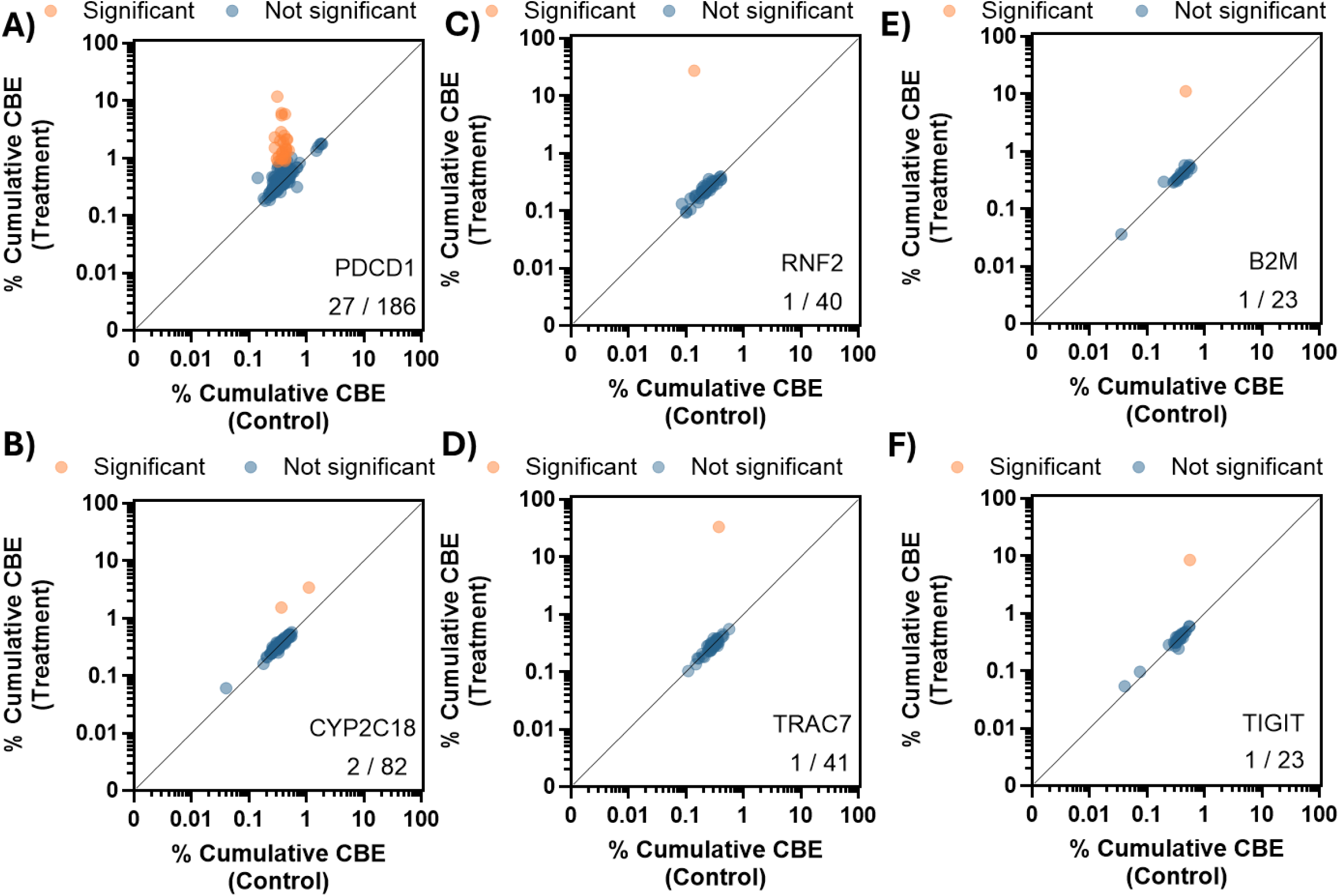
Confirmation in HSPCs with wildtype *S.p.* Cas9-CBE delivered as mRNA (CBE). Editing quantification at assays with >1,000x coverage. Each dot represents a gRNA on/off-target with the average raw frequency of cumulative CBE transition events of the control (x-axis) and treatment (y-axis) plotted. Blue dots indicate sites with no statistical significance while orange dots indicate significant sites (p adj<0.05) for the **A)** *PDCD1* site 8, **B)** *CYP2C18*, **C)** *RNF2*, **D)** T*RAC* site 7, **E)** *B2M* site 1, and **F)** *TIGIT* site 7 gRNAs. Text at the bottom right displays the total number of confirmed sites out of all interrogated with sufficient depth (>1,000x).

**Supplementary Figure 13.**
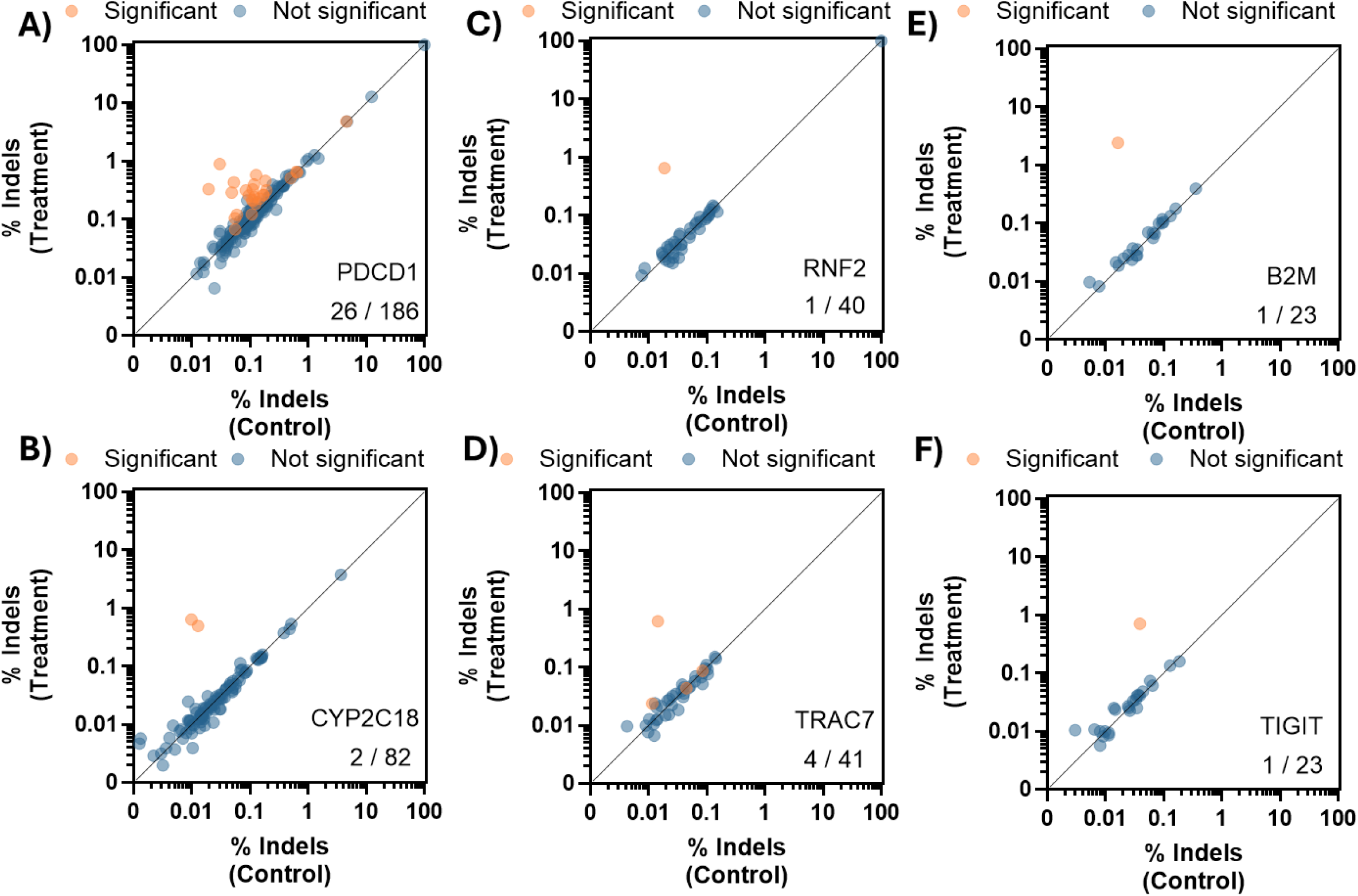
Confirmation in HSPCs with wildtype *S.p.* Cas9-ABE delivered as mRNA (indels). Editing quantification at assays with >1,000x coverage. Each dot represents a gRNA on/off-target with the average raw frequency of indel events of the control (x-axis) and treatment (y-axis) plotted. Blue dots indicate sites with no statistical significance while orange dots indicate significant sites (p adj<0.05) for the **A)** *PDCD1* site 8, **B)** *CYP2C18*, **C)** *RNF2*, **D)** T*RAC* site 7, **E)** *B2M* site 1, and **F)** *TIGIT* site 7 gRNAs. Text at the bottom right displays the total number of confirmed sites out of all interrogated with sufficient depth (>1,000x).

**Supplementary Figure 14.**
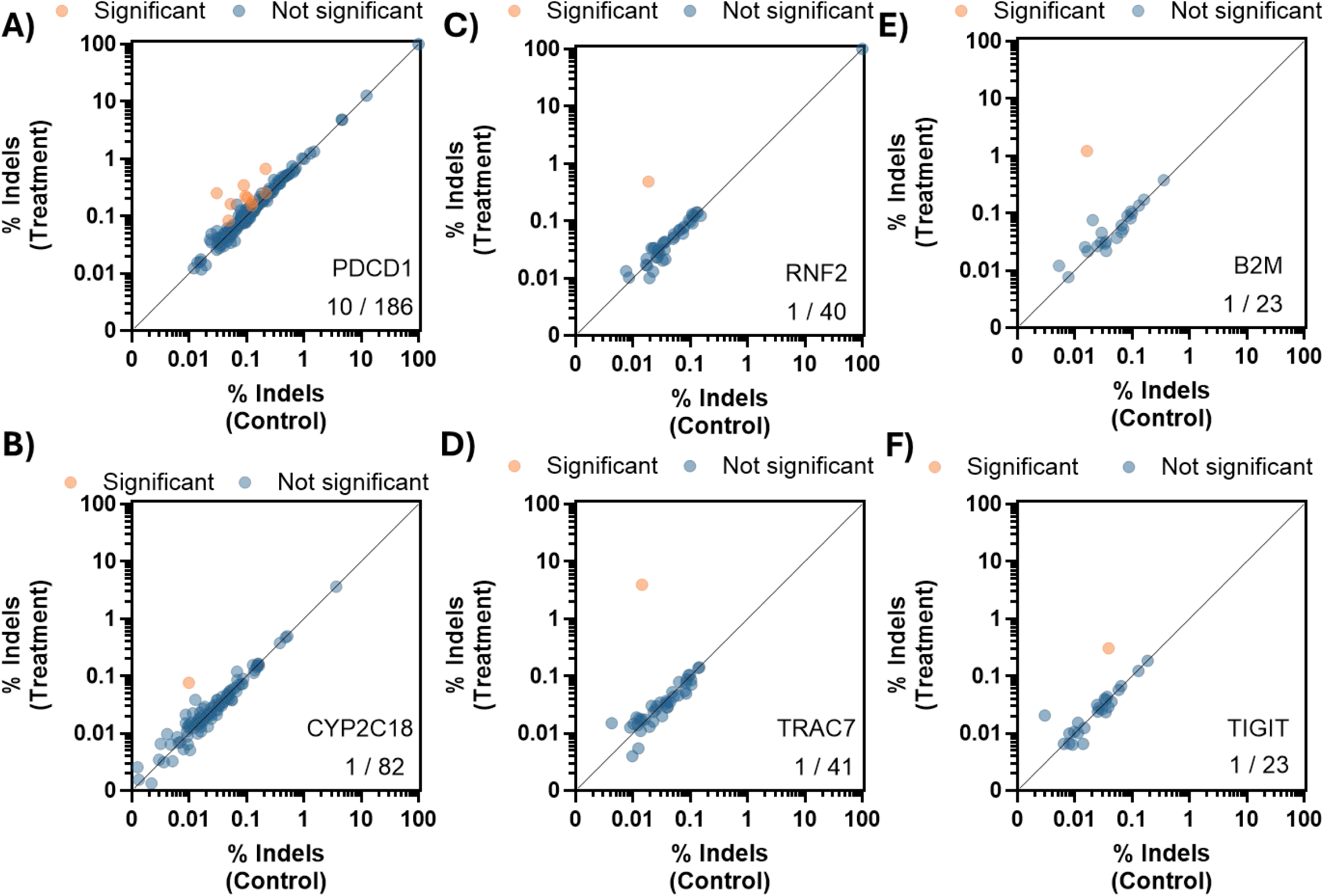
Confirmation in HSPCs with wildtype *S.p.* Cas9-CBE delivered as mRNA (indels). Editing quantification at assays with >1,000x coverage. Each dot represents a gRNA on/off-target with the average raw frequency of indels of the control (x-axis) and treatment (y-axis) plotted. Blue dots indicate sites with no statistical significance while orange dots indicate significant sites (p adj<0.05) for the **A)** *PDCD1* site 8, **B)** *CYP2C18*, **C)** *RNF2*, **D)** T*RAC* site 7, **E)** *B2M* site 1, and **F)** *TIGIT* site 7 gRNAs. Text at the bottom right displays the total number of confirmed sites out of all interrogated with sufficient depth (>1,000x).

**Supplementary Figure 15.**
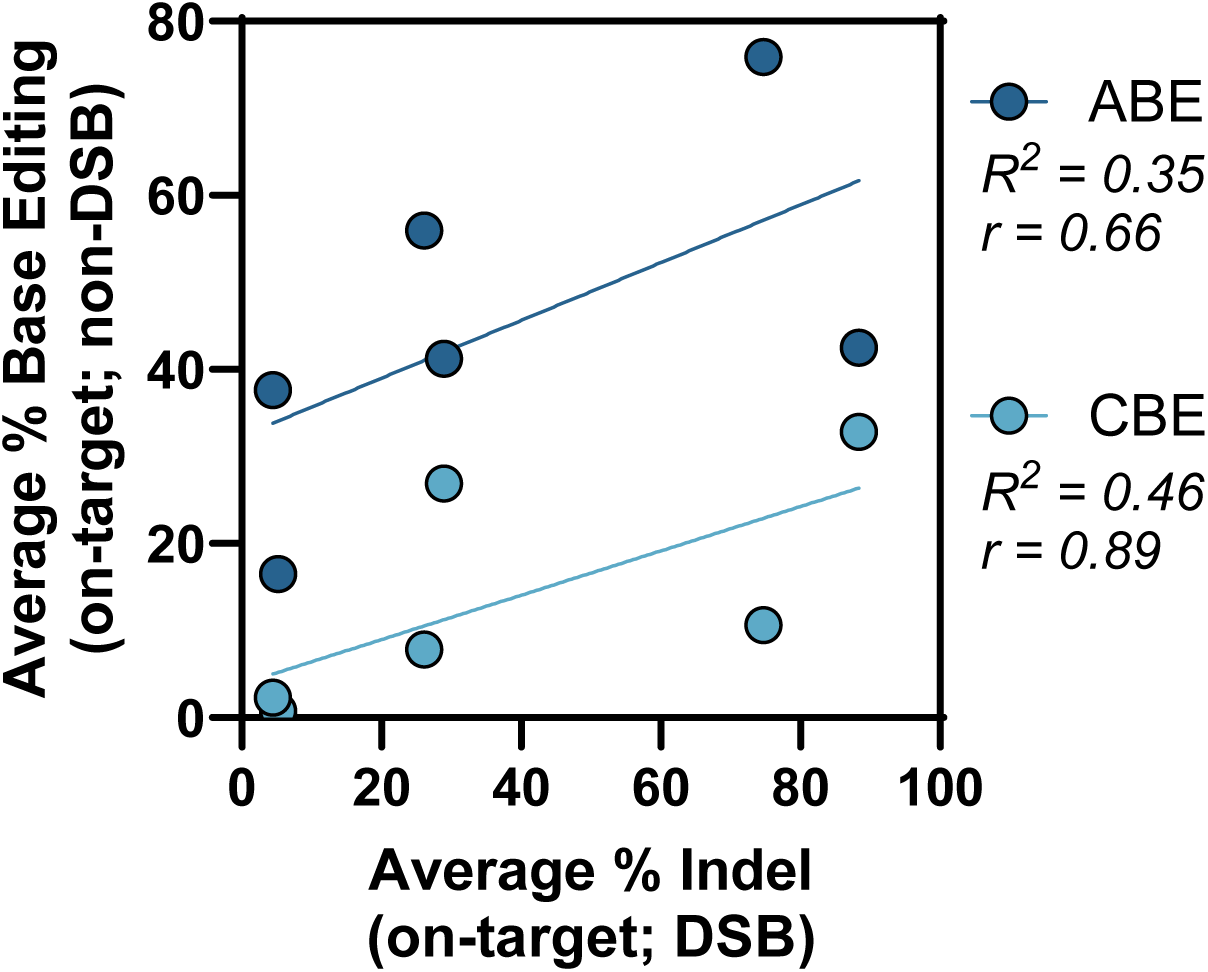
On-target indel and base editing correlations in HSPCs. Frequencies were plotted for all on-target ABE/CBE sites with both SSB and DSB along with the respective DSB indel frequency (wildtype Cas9) and Spearman r calculated.

